# OCA-B/Pou2af1 Expression in T Cells Promotes PD-1 Blockade-Induced Autoimmunity but is Dispensable for Anti-Tumor Immunity

**DOI:** 10.1101/2025.10.22.683978

**Authors:** Junhong Du, Asit K. Manna, Miguel A. Medina-Serpas, Erik P. Hughes, Paulla Bisoma, Kimberley J. Evason, Arabella Young, W. David Wilson, Todd M. Brusko, Abdelbasset A. Farahat, Dean Tantin

## Abstract

The transcription coregulator OCA-B promotes CD4^+^ T cell memory recall responses and autoimmunity. OCA-B T cell deletion prevents spontaneous type-1 diabetes (T1D) onset in non-obese diabetic (NOD) mice and blunts T1D in a subset of more aggressive models. However, the role of OCA-B in diabetes induced by treatment with immune checkpoint inhibitors (ICIs), and the role of OCA-B in the control of tumors with and without ICI treatment, has not been studied. Here we show that islet and pancreatic lymph node T cells from T1D individuals express measurable *POU2AF1* mRNA. Deletion of OCA-B in T cells fully insulates 8-week-old non-obese diabetic (NOD) mice against ICI-induced diabetes and partially protects 12-week-old mice. Salivary and lacrimal gland infiltration and inflammation were also reduced. Protection was associated with a block in the differentiation of progenitor exhausted CD8^+^ T cells (T_PEX_) into terminally exhausted CD8^+^ T cells (T_EX_). We show that OCA-B T cell loss preserves anti-tumor immune responses following PD-1 blockade in different tumors and mouse strains. These findings point to a potential therapeutic window in which pharmaceuticals targeting OCA-B could be used to block the emergence of both spontaneous and ICI-induced autoimmunity while sparing anti-tumor immunity. We develop first-in-class small molecule inhibitors of Oct1/OCA-B transcription complexes and show that administration into NOD mice also blocks diabetes emergence following PD-1 blockade. These results identify OCA-B as a promising therapeutic target for the prevention of autoimmunity and immune-related adverse events (irAEs).

## INTRODUCTION

Type 1 diabetes (T1D) affects 1.8 million people in the United States and incidence is increasing worldwide (*1–3*). It is characterized by autoimmune destruction of pancreatic β cells and consequent inability to produce insulin (*4*). While T1D symptoms can be managed with insulin injections, they must be taken for life. Transplantation of islets of Langerhans can restore insulin-independence to T1D patients, but results in reactivation of islet-specific memory cells T cells, β cell destruction and transplant rejection (*5, 6*). Similarly, engraftment of islets from non-obese diabetic (NOD) mice with severe combined immunodeficiency (NOD.SCID) into diabetic NODs briefly restores normoglycemia, but results in robust memory autoimmune responses and complete loss of grafted islet function in 10-14 days (*7*). Human therapies aim to minimize graft rejection or block the initial emergence of T1D in at-risk patients altogether while sparing normal immune function (*8*).

In humans, α-PD-1 therapy as well as therapies directed against a number of other inhibitory T cell receptors (immune checkpoint inhibitors or ICIs) are now used for treatment of a variety of cancer indications (*9–12*). The development and wide adoption of ICIs has resulted in a concomitant increase in associated unwanted immune-related adverse events, or irAEs, affecting a variety of tissues (e.g., skin, rheumatoid, pulmonary, gut, cardiac), some of which can be permanent (*10, 13–21*). IrAEs include endocrinopathies such as thyroiditis (*22*), and less frequently hypophysitis, autoimmune diabetes and other diseases (*17, 19, 21, 23*). A critical gap in the field is the identification of treatment strategies that block the emergence of irAEs while preserving anti-tumor immunity.

Polymorphisms in the *HLA-DR* and *HLA-DQ* genes encoding MHC class II have the strongest genetic influence on T1D, accounting for 40-50% of familial risk (*24*). These findings implicate CD4^+^ T cells as primary drivers of pathogenesis. CD4^+^ T cells direct other cell types such as CD8^+^ T cells and macrophages to infiltrate pancreatic islets and cause β-cell destruction (*25*). Consistent with these findings, transient T cell depletion using α-CD3ε antibodies (Teplizumab-mzwv) significantly delays T1D onset in at-risk individuals in human clinical trials (*26–29*). Though promising, such treatments are limited by the resulting global immunosuppression. An ideal alternative would selectively deplete pathogenic T cell function while preserving beneficial immune function, including immunity against tumors.

OCA-B is a B cell- and T cell-restricted transcription coactivator (*30, 31*). In CD4^+^ T cells, OCA-B docks with the transcription factor Oct1 (gene symbol *Pou2f1*) or Oct2/Pou2f2 to regulate targets including *Il2*, *Ifng*, *Icos*, *Csf2* and *Tbx21* (*32, 33*). Unlike transcription factors such as NFAT, AP-1 and NF-κB, OCA-B controls the expression of these genes specifically during repeated antigen exposures (*32, 33*). Agents targeting this pathway would selectively inhibit over-active T cell responses to antigen re-exposure but preserve acute immune response. Consistent with this prediction, in the absence of either OCA-B or Oct1 primary T cell responses to systemic and glial-tropic viruses (LCMV and JHMV, respectively) and viral clearance are normal (*31, 34*), however fewer central memory (T_CM_) CD4^+^ T cells form, and those that do are highly deficient in their ability to respond to antigenic re-challenge (*31, 33*). Three studies directly link polymorphisms in the human *OCAB* (*POU2AF1*) locus with autoimmune disease (*35–37*). Human polymorphisms affecting Oct1/2 (and therefore OCA-B) binding to the *TNFA* and *CTLA4* loci are associated with inflammatory bowel disease and systemic lupus erythematosus, respectively (*38, 39*). Additionally, two meta-analyses show a strong association between Oct1/Oct2/OCA-B binding site polymorphisms and autoimmunity including T1D(*40, 41*). Collectively, these studies link OCA-B with human autoimmunity.

OCA-B loss in T cells protects mice against spontaneous T1D onset (*42*). Single-cell RNA-sequencing (scRNA-seq) reveals that protection is associated with loss of autoreactive CD8^+^ T cell clones but maintenance of the overall pancreatic islet immune architecture (*42*). These findings suggest that targeting OCA-B could be used to treat T1D early in the disease course while sparing baseline immunity. OCA-B’s lymphoid-restricted expression pattern (*30, 31*), its dispensability for primary antiviral responses (*31, 33, 34*), and its specificity for Oct1/Pou2f1 and Oct2/Pou2f2 complexes with DNA (*43–45*) make it a promising therapeutic target that could be used to specifically block autoreactive CD4^+^ T cells while sparing normal immunity. OCA-B peptide mimics linked to membrane-penetrating peptides can reduce inflammation and hyperglycemia in recent-onset T1D NOD mice, but also show significant toxicity (*42*). An outstanding question is whether OCA-B can be successfully targeted by more drug-like small molecules. It also remains to be determined if OCA-B promotes diabetes driven by ICI therapy, and/or if OCA-B promotes anti-tumor immunity, including that promoted by ICI.

Here, we identify T cells expressing POU2AF1 in the pancreatic lymph nodes and islets of T cells expressing inflammatory markers in individuals with early T1D. We show that OCA-B in T cells promotes the emergence of autoimmune diabetes following ICI administration in NOD mice, while being dispensable for anti-tumor immune responses following ICI administration. We develop a small molecule DNA-binding inhibitor that effectively disrupts Oct1:OCA-B complex in vitro and OCA-B’s transcription functions in cultured cells. We show that administration of these compounds to NOD mice blocks the emergence of diabetes and prevents inflammation in pancreatic islets and other endocrine organs following ICI administration, with in vivo efficacy tracking with activity in vitro. Collectively, the findings indicate that OCA-B promotes diabetes development following ICI treatment, and that targeting OCA-B could be used to block irAEs following ICI while sparing anti-tumor immunity.

## RESULTS

### Detection of POU2AF1/OCAB mRNA in proinflammatory T cells in the setting of early human T1D

T cell conditional OCA-B loss protects NOD mice against spontaneous T1D (*42*). To identify a possible association with T cells in the context of human T1D, we performed a targeted reanalysis of previously published spatial transcriptomic data (*46*) using collated transcriptomic information from an individual with T1D (nPOD6551, 20 years old, 0.58 years duration) to identify CD8^+^ T cells (pseudocolored yellow), B cells (green) and islet cells (white), as well as targeted transcript annotations for *CHGA* (pseudocolored red) to highlight β cells, *CD8A* (purple) to highlight T cells, *MS4A1* (orange) to highlight B cells. *POU2AF1* was pseudocolored cyan. Focusing on a representative islet with obvious insulitis, we observed strong peri-islet inflammation with both T cells and rare B cells (**Fig. 1A**). The B cells in the field all displayed obvious POU2AF1 gene expression as a positive control, while approximately 9.4% of CD8^+^ T cells (19/202 cells) showed measurable *POU2AF1* expression. *POU2AF1*-expressing CD4^+^ and CD8^+^ T cells were also identified in the pancreatic lymph nodes (pLNs) from the same individual (**Fig. 1B-C**). These findings confirm the presence of *POU2AF1*-expressing T cells within the pancreas of T1D patients.

**Fig. 1.**
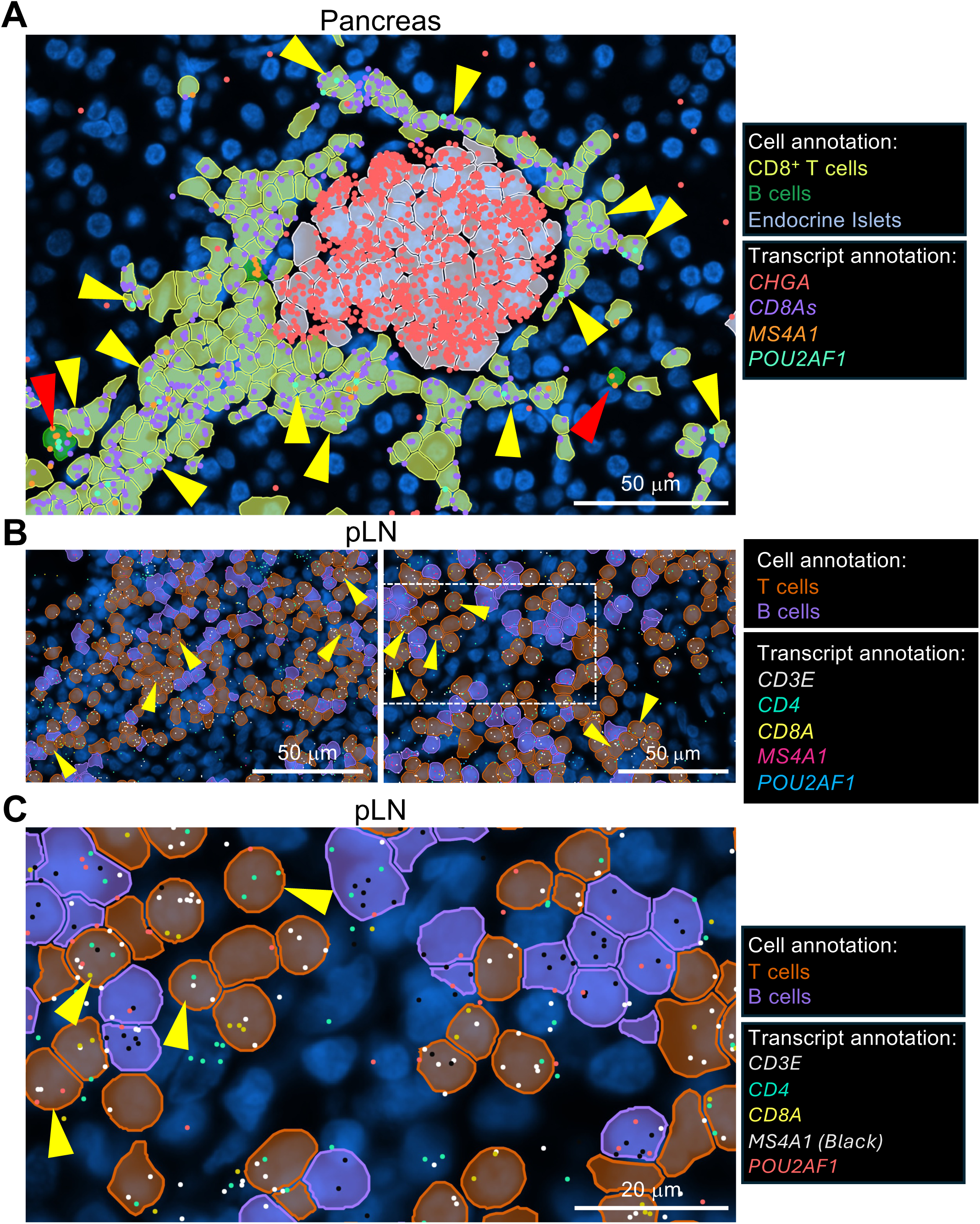
POU2AF1 labels pathogenic T cells in the setting of human T1D. **(A)** 10X Xenium image of a pancreatic islet from a 20-year-old human patient with early T1D. *CHGA* (red), *CD8A* (purple), *MSFA1* (orange) and *POU2AF1* (green) transcripts are annotated. Cells were identified by imputation as CD8+ T cells (yellow), B cells (green), or islet endocrine cells (gray). Yellow arrows highlight *POU2AF1*-expressing CD4^+^ T cells and red arrows highlight *POU2AF1*-expressing B cells. (**B**) Data from pLN sections of the same individual. *CD3E* (white), *CD4* (green), *CD8A* (yellow), *MS4A1* (magenta), *POU2AF1* (blue). Yellow arrows highlight T cells that also express *POU2AF1*. (**C**) Detail from the white box in the right panel shown in (B), with a slightly different transcript annotation color scheme: *CD3E* (white), *CD4* (green), *CD8A* (yellow), *MS4A1* (black), *POU2AF1* (red). B cells expressing *POU2AF1* were abundant and are not highlighted.

### OCA-B T cell knockout protects NOD mice against diabetes induced by PD-1 blockade

Modulating the PD-1 pathway induces diabetes and other endocrinopathies in NOD mice (*47–50*). To test the effect of OCA-B T cell loss in diabetes induced by PD-1 blockade, we treated 8-week-old male NOD mice with four injections of α-PD-1 antibody to follow the disease timecourse, or one injection of α-PD-1 antibody with a 5-day interval prior to flow cytometric analysis of NOD mice (**Fig. 2A**). OCA-B loss in T cells completely blocked α-PD-1-induced diabetes development (**Fig. 2B-C**). Treatment of 12-week-old female NOD mice with 5 total injections of α-PD-1 resulted in less complete protection, with half the mice failing to develop diabetes and the other half showing delayed onset (**fig. S1A-B**). Flow cytometry revealed a significant reduction in islet-infiltrating CD45^+^ leukocytes in α-PD-1-treated knockout mice compared to littermate controls (**Fig. 2D-E**). We also observed a decrease in the ratio of islet-infiltrating CD8^+^/CD4^+^ T cells in 8-week-old knockouts (**Fig. 2F-G**). Autoreactive CD8^+^ T cells specifically recognize and destroy β cells and their relative numbers tend to increase with PD-1 blockade (*25*). Among the infiltrating CD8^+^ cells, there were no changes in total CD11a^+^CD62L^lo^ activated cells (**Fig. 2H-I**), suggesting that OCA-B does not regulate acute activation in this context, however CD8^+^PD-1^+^ cells were significantly reduced (**Fig. 2J-K and fig. S1C-D**). PD-1 blockade results in the proliferation and differentiation of exhausted CD8^+^ T cells in pancreatic islets, promoting the expansion of both T_PEX_ and T_EX_ cells but also skewing the islet total exhausted T cell population in favor of T_EX_ (*25*). To determine the effect of OCA-B loss on this pathogenic maturation, we used SLAMF6 (also known as Ly108) and CD39 to distinguish CD8^+^ T_PEX_ and T_EX_, respectively, at both 8 and 12 weeks. OCA-B loss resulted in a marked diminution in T_EX_ percentages among both PD-1^+^CD8^+^ and total islet cells (**Fig. 2L-M**), consistent with a defect in the ability of CD8^+^ T_PEX_ cells within the islets of 8-week-old mice to differentiate into T_EX_. Similar findings were made at 12 weeks though with decreased magnitude (**fig. S1E-F**). A similar decrease of SLAMF6^-^CD39^+^ terminally exhausted CD4^+^ T cells was also observed (**fig. S1G-H**). In the absence of α-PD-1 treatment, T_PEX_ percentages were unaffected while knockouts still showed a substantial decrease in T_EX_, suggesting that the differences in T_EX_ with α-PD-1 treatment are not due to prior differences in T_PEX_ abundance in the knockouts (**fig. S2A-B**). Lastly, following PD-1 blockade in 8-week-old animals, we observed stark differences in the accumulation of CD62L^hi^ T_PEX_, an even more developmentally plastic CD8^+^subpopulation (*51*) (**fig. S2C-D**). Cumulatively, these results indicate that OCA-B loss in T cells blocks the induction of diabetes by α-PD-1 treatment and reduces the pathogenic maturation of CD8^+^ T_PEX_ in the pancreatic islets.

**Fig. 2.**
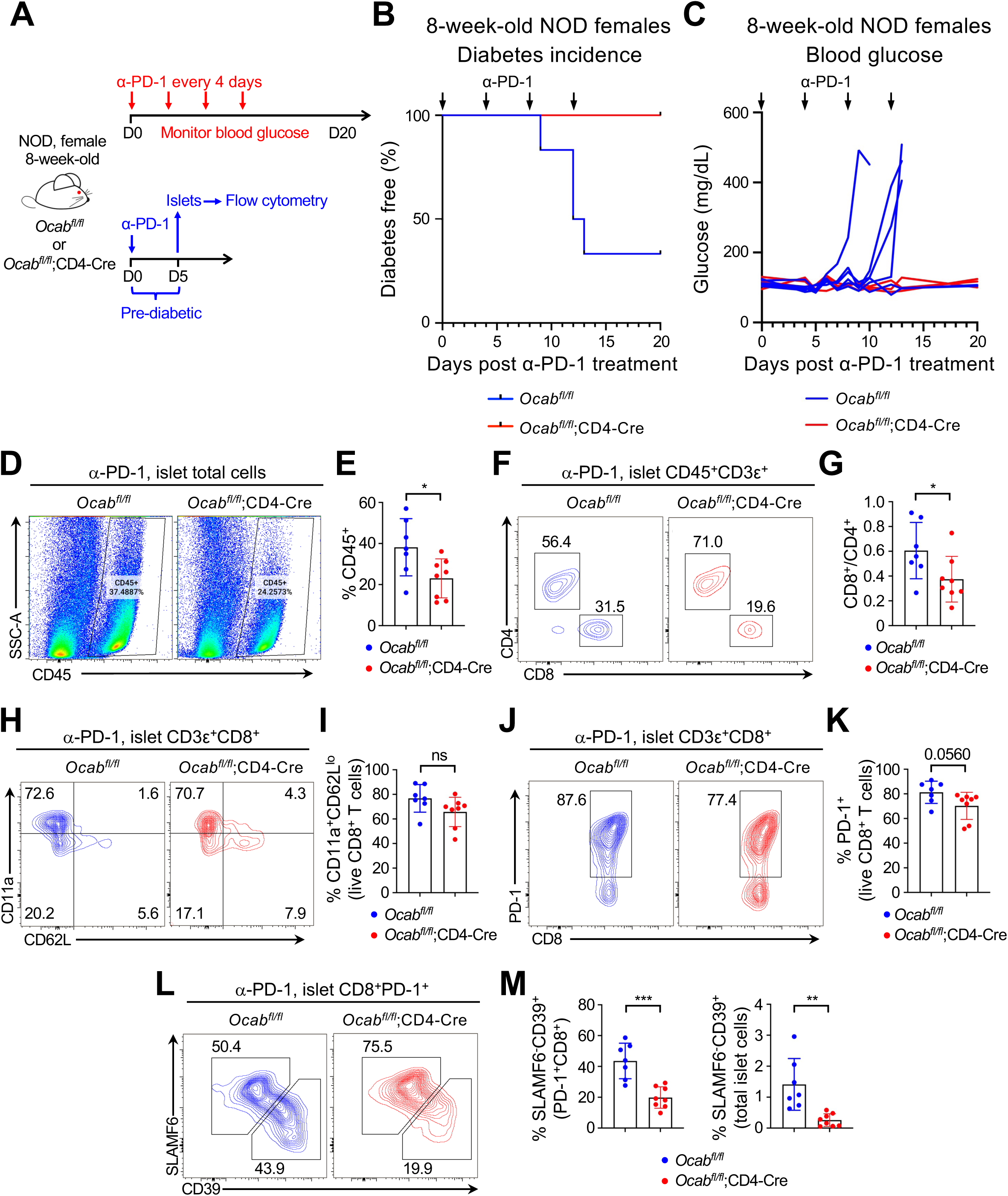
OCA-B deletion in T cells confers protection against PD-1 blockade-induced diabetes in NOD mice. **(A)** Schematics of experimental workflow for monitoring α-PD-1-induced autoimmune diabetes in 8-week-old NOD male *Ocab^fl/fl^* and *Ocab^fl/fl^*;CD4-Cre injected with α-PD-1, and monitoring pre-diabetic mice at short timepoints (day 5) after a single α-PD-1 injection. **(B)** Kaplan-Meier plot of diabetes-free 8-week-old NOD male *Ocab^fl/fl^* (n=7) and *Ocab^fl/fl^*;CD4-Cre (n=7) mice. 250 µg α-PD-1 was administered by *i.p.* injection on day 0, 4, 8 and 12. Blood glucose levels were measured every 1-2 days, with mice considered diabetic if glucose rose above 250 mg/dL over two consecutive days. **(C)** Blood glucose levels of individual mice from (B). **(D)** Islets were prepared from 8-week-old treated knockout and littermate control female mice 5 days after one dose of α-PD-1 antibody for flow cytometry. Representative flow cytometry plots depicting total infiltrating CD45^+^ leukocytes for control and knockout mice. (**E**) Quantification of total infiltrating CD45^+^ islet leukocyte flow cytometry data from NOD.*Ocab^fl/fl^* (n=7) and NOD.*Ocab^fl/fl^*;CD4-Cre mice (n=8). (**F**) Representative flow cytometry plots depicting CD4^+^ and CD8^+^ islet T cells. Cells were pre-gated on CD45^+^CD3ε^+^. (**G**) Quantification of CD8^+^/CD4^+^ ratio from NOD.*Ocab^fl/fl^* (n=7) and NOD.*Ocab^fl/fl^*;CD4-Cre mice (n=8). (**H**) Representative flow cytometry plots showing CD62L and CD11a expressions of islet-infiltrating CD8^+^ T cells. (**I**) Quantification of activated CD62L^-^CD11a^+^CD8^+^ T cells from NOD.*Ocab^fl/fl^*(n=7) and NOD.*Ocab^fl/fl^*;CD4-Cre mice (n=8). (**J**) Representative flow cytometry plots depicting PD-1-expressing CD8^+^ islet-infiltrating T cells. Cells were pre-gated on CD3ε^+^CD8^+^. (**K**) Quantification of the percentage of PD-1^+^ T cells among CD8^+^ T cells from NOD.*Ocab^fl/fl^* (n=7) and NOD.*Ocab^fl/fl^*;CD4-Cre mice (n=8). (**L**) Representative flow cytometry plots depicting the percentage of islet SLAMF6^+^CD39^-^T_PEX_ and SLAMF6^-^CD39^+^ T_EX_. Cells were pre-gated on PD-1^+^CD8^+^. (**M**) Quantification of flow cytometry data for the percentages of T_EX_ among CD8^+^PD-1^+^ and total cells in islets from NOD.*Ocab^fl/fl^* (n=7) and NOD.*Ocab^fl/fl^*;CD4-Cre mice (n=8).

### OCA-B T cell knockout maintains anti-tumor immunity induced by PD-1 blockade

To determine if OCA-B T cell loss has a corresponding effect on anti-tumor immunity, we used C57BL/6 and NOD mice OCA-B conditional knockout mice (*31, 42*) with syngeneic tumor lines. In the C57BL/6 strain background we used two lines, B16F10 and MC38. Compared to littermate controls lacking the CD4-Cre driver, OCA-B loss had no effect on B16F10 tumor growth (**fig. S3**). This result was expected because unmodified B16 cells do no elicit robust immune responses unless given a combination of ICI treatment (*52*). In contrast, “immune-hot” MC38 colon adenocarcinoma cells exhibit significant control by T cells that can be augmented by α-PD-1 treatment (*53*). We found that MC38 tumor control was preserved with OCA-B T cell loss either in the absence or presence of α-PD-1 treatment (**Fig. 3A-B**). Flow cytometry of cells from the tumor microenvironment (TME) showed equivalent CD45.2^+^ leukocyte infiltration and CD8^+^ to CD4^+^ T cell ratios with α-PD-1 treatment (**Fig. 3C-D**). The percentages of CD4^+^ and CD8^+^ SLAMF6^+^CD39^-^ and SLAMF6^-^CD39^+^ cells in the TME following α-PD-1 antibody treatment were similar between OCA-B knockouts and controls (**Fig. 3E-F**). T cell exhaustion in tumor-infiltrating CD4^+^ and CD8^+^ as indicated by TIM-3 expression was also equivalent (**Fig. 3G-H**).

**Fig. 3.**
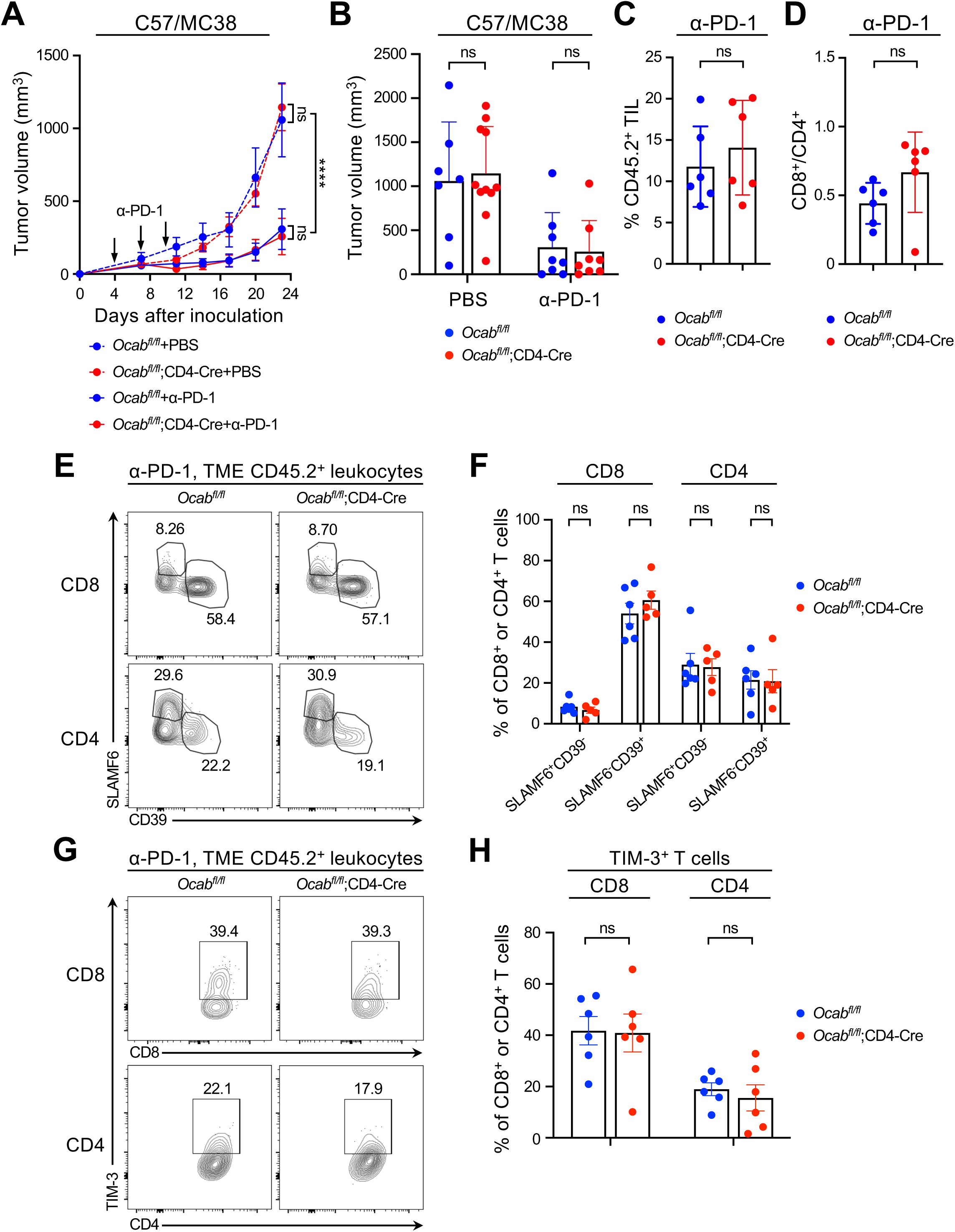
OCA-B in T cells is dispensable for anti-tumor immunity in C57BL/6 mice. **(A)** 5×10^5^ MC38 tumor cells were *s.c*. implanted into the hind flanks of female C57BL/6 *Ocab^fl/fl^* and *Ocab^fl/fl^*;CD4-Cre mice. 300 µg α-PD-1 (or PBS) was administered *i.p.* on day 4, 7 and 10. Tumor sizes were measured every 3-4 days, and mice were euthanized at end point (day 23) or upon tumor size reaching 2 cm in any direction. *Ocab^fl/fl^*+PBS (n=7), *Ocab^fl/fl^*;CD4-Cre+PBS (n=11), *Ocab^fl/fl^*+α-PD-1 (n=8), and *Ocab^fl/fl^*;CD4-Cre+α-PD-1 (n=8). **(B)** Mean tumor sizes from (A) are shown. *Ocab^fl/fl^*+PBS (n=7), *Ocab^fl/fl^*;CD4-Cre+PBS (n=11), *Ocab^fl/fl^*+α-PD-1 (n=8), and *Ocab^fl/fl^*;CD4-Cre+α-PD-1 (n=8). **(C)** Quantification of flow cytometry data showing the percentage of CD45.2^+^ tumor-infiltrating leukocytes (TILs) in MC38 TMEs from α-PD-1 antibody-treated *Ocab^fl/fl^*(n=6) and *Ocab^fl/fl^*;CD4-Cre (n=6) mice on day 23 or upon tumor size reaching 2 cm in any direction. **(D)** Quantification of flow cytometry data showing the ratio of CD8^+^ to CD4^+^ T cells in the TME. **(E)** Representative flow cytometry plots of endpoint TME CD4^+^ and CD8^+^ T cell SLAMF6 and CD39 expression. *Ocab^fl/fl^* (n=6) and *Ocab^fl/fl^*;CD4-Cre (n=5) tumor-bearing mice were treated with α-PD-1 antibodies. (**F**) Quantification of flow cytometry data for endpoint TME CD4^+^ and CD8^+^ T cell SLAMF6 and CD39 expression from *Ocab^fl/fl^* (n=6) and *Ocab^fl/fl^*;CD4-Cre (n=5) mice. (**G**) Representative flow cytometry data for endpoint TME CD8^+^ and CD4^+^ T cell TIM-3 expression. (**H**) Quantification of flow cytometry data at endpoint showing TME CD4^+^ and CD8^+^ T cell TIM-3 expression from *Ocab^fl/fl^* (n=6) and *Ocab^fl/fl^*;CD4-Cre (n=6) mice.

Because the protection from PD-1 blockade-induced diabetes was observed using NOD background mice, we also assessed tumor responses using a syngeneic “immune-hot” 9495 sarcoma cell line (*54*), allowing us to measure diabetes incidence and tumor control in the same animals. α-PD-1 antibody treatment was effective in controlling 9495 tumor growth in syngeneic NOD mice (**Fig. 4A**) as expected (*54*). Moreover, tumor control either with or without α-PD-1 treatment was equivalent regardless of OCA-B status (**Fig. 4A**). A similar cohort of mice were again fully protected against diabetes in the OCA-B deficient group, though penetrance in the control group was lower due to the decreased amount of α-PD-1 antibody used (200 μg vs. 250 μg) to slow diabetic progression and allow longer monitoring of tumor growth (**fig. S4A**). Although statistically nonsignificant, there was a trending decrease in CD45^+^ leukocyte tumor infiltration in OCA-B-deficient mice (**Fig. 4B**). This was driven by 4 control animals with high tumor infiltrate levels. As with MC38 tumors in C57BL/6 mice, CD8^+^ percentages, T_PEX_ to T_EX_ differentiation, and CD8^+^ TIM-3 expression (**Fig. 4C-H**) were equivalent in the context of α-PD-1 treatment. There were also few differences in CD4^+^ and CD8^+^ T cell activation and PD-1 expression (**fig. S4B-E**). These findings indicate minimal differences in anti-tumor immune responses to multiple syngeneic tumor lines in OCA-B T cell knockout mice, indicating a divergence between anti-tumor immunity and the precipitation of irAEs that could be leveraged therapeutically.

**Fig. 4.**
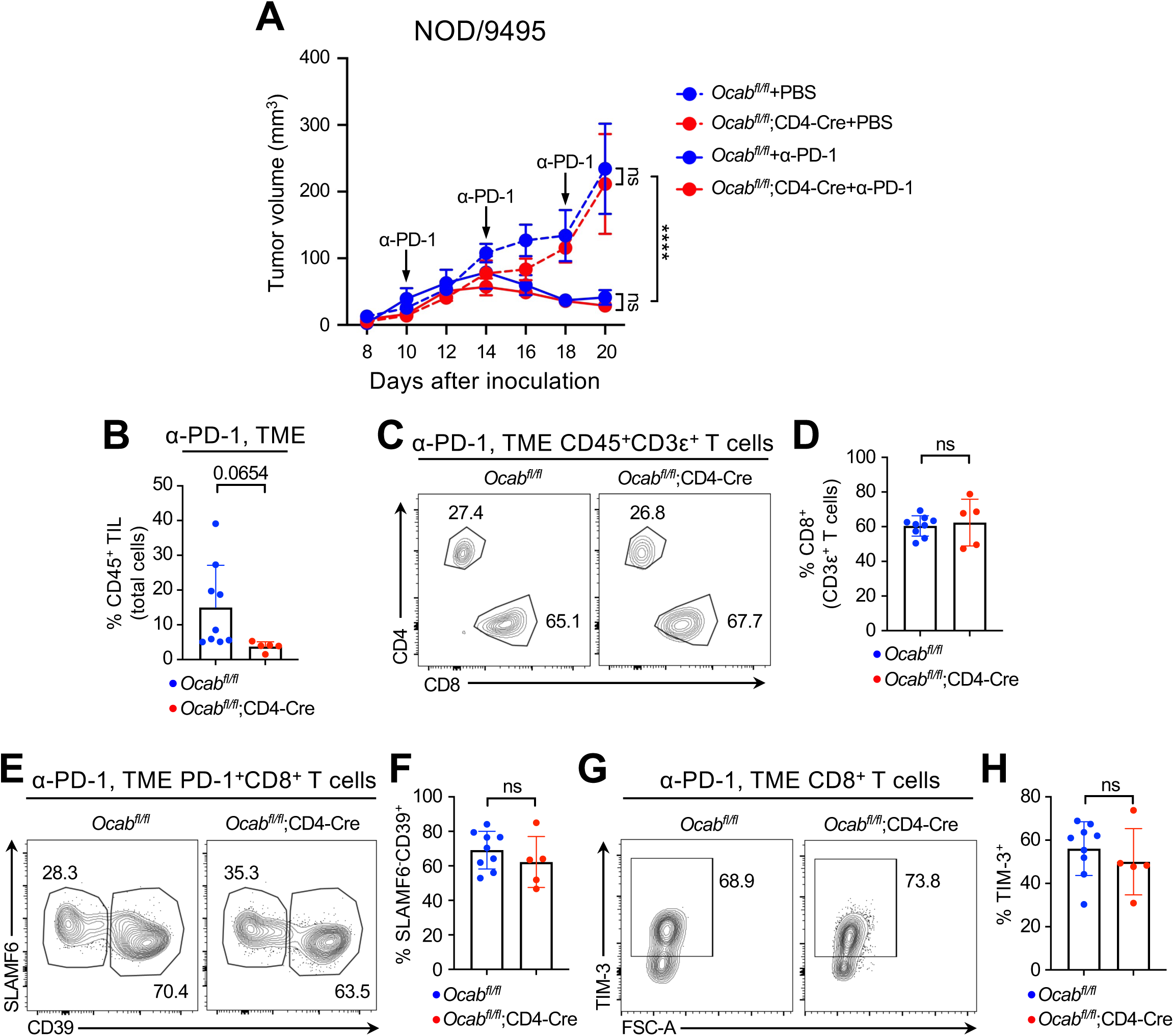
OCA-B in T cells is dispensable for anti-tumor immunity in NOD mice. **(A)** 1.5×10^6^ 9495 tumor cells were *s.c*. implanted into the hind flanks of male NOD.*Ocab^fl/fl^*and NOD.*Ocab^fl/fl^*;CD4-Cre mice. Three doses of 200 µg α-PD-1 (or PBS vehicle control) were *i.p.* administered on day 10, 14 and 18. Tumor sizes were measured during different stages of tumor growth. *Ocab^fl/fl^*+PBS (n=6), *Ocab^fl/fl^*;CD4-Cre+PBS (n=7), *Ocab^fl/fl^*+α-PD-1 (n=8), and *Ocab^fl/fl^*;CD4-Cre+α-PD-1 (n=12). **(B)** Quantification of flow cytometry data for the percentage of CD45^+^ infiltrating cells in the TME from α-PD-1 antibody-treated control *Ocab^fl/fl^* (n=9) and littermate experimental *Ocab^fl/fl^*;CD4-Cre (n=5) mice. **(C)** Representative flow cytometry plots depicting CD4^+^ and CD8^+^ TME T cells from α-PD-1 antibody-treated control NOD.*Ocab^fl/fl^* and littermate experimental NOD.*Ocab^fl/fl^*;CD4-Cre mice. Cells were pre-gated using α-CD3 staining. (B) Quantification of flow cytometry data depicting the percentage of CD4^+^ and CD8^+^ T cells in the TME from NOD.*Ocab^fl/fl^* (n=9) and NOD.*Ocab^fl/fl^*;CD4-Cre mice (n=5). (**E**) Representative flow cytometry plots depicting relative percentages of SLAMF6^-^ and CD39^-^expressing infiltrating CD8^+^PD-1^+^ T cells. (**F**) Quantification of PD-1^+^SLAMF6^+^CD39^-^CD8^+^ T_PEX_ and PD-1^+^SLAMF6^-^CD39^+^CD8^+^ T_EX_ among PD-1^+^CD8^+^ T cells in the TME from NOD.*Ocab^fl/fl^* (n=9) and NOD.*Ocab^fl/fl^*;CD4-Cre mice (n=5). (**G**) Representative flow cytometry plots depicting TIM-3 levels in infiltrating CD8^+^ T cells. (**H**) Quantification of TIM-3^+^ cells among CD8^+^ T cells in the TME from NOD.*Ocab^fl/fl^* (n=9) and NOD.*Ocab^fl/fl^*;CD4-Cre mice (n=5). In (panels C-H), one data point from NOD.*Ocab^fl/fl^* was removed because of low cell numbers.

### Development of small molecule inhibitors of Oct1:OCA-B complexes

The Oct1:OCA-B:DNA co-crystal structure (*45*) suggests multiple possible targeting strategies, including rationally designed OCA-B peptide mimics (**Fig. 5A**, upper lightning bolt) which are functional but associated with toxicity in cells and mice due to the membrane-penetrating peptide (*42*). To identify novel small molecule inhibitors, we used another strategy in which DNA minor groove-binding compounds with specificity for the consensus octamer motif (5’ATGCAAAT) would disrupt Oct1:OCA-B complexes. These inhibitors would dock on the underside of the DNA in the structure shown in **Fig. 5A** (bottom lightning bolt) and would interfere with the unstructured Oct1 linker domain that also spans the minor groove. We synthesized and tested 9 candidate minor groove-binding compounds with specificity for the octamer motif. To test efficacy, we developed two cell-based assay systems. The first is a transcription-based platform in which we stably transfected the human T cell line SupT1 to express secreted nanoluciferase (nLuc) under the control of five Oct1/OCA-B binding sites. SupT1 T cells express OCA-B at higher levels than, e.g. Jurkat cells, and at levels comparable to primary thymocytes (**Fig. 5B**). We generated a stable-transfected SupT1 clone expressing nLuc under the control of five wild-type octamer sequences (“5×Oct”) and a second using mutant sites incapable of binding Oct1 and OCA-B (“5×Mut”, **Fig. 5C**). The 5×Oct clone produced substantial levels of luciferase activity relative to empty vector-transduced controls, while the 5×Mut clone expressed lower but still measurable activity (**Fig. 5D**). Timecourse experiments show that 12- to 18-hr incubations have not yet saturated nLuc activity and capture a dynamic range allowing both activators and inhibitors to be detected (**Fig. 5E**). We therefore chose 18-20 hr for all further experiments. Cell counts were also taken to measure cell viability, using 2.5 μM staurosporine a positive control for maximum toxicity. The treatment kills >98% of SupT1 cells within ∼20 hr and correspondingly inhibits luciferase expression (**fig. S5**). Of the 9 candidate compounds two – DB1033 (IC_50_∼26 μM, **Fig. 5F**) and DB2232 (IC_50_∼10 μM, **fig. S6A**) – inhibited luciferase activity without significantly affecting luciferase expression from the mutant cell line and without significant toxicity. Other compounds such as DB2658 showed significant toxicity in the same screen (**fig. S6B**) or were less active (e.g., DB2385, **fig. S6C-H**). The DB1033 structure is shown in **Fig. 5G**. Molecular docking indicated that DB1033 preferentially interacts with DNA minor groove in the AAAT portion of the consensus octamer sequence, overlapping with the location of the unstructured Oct1 linker domain (**Fig. 5H-I**).

**Fig. 5.**
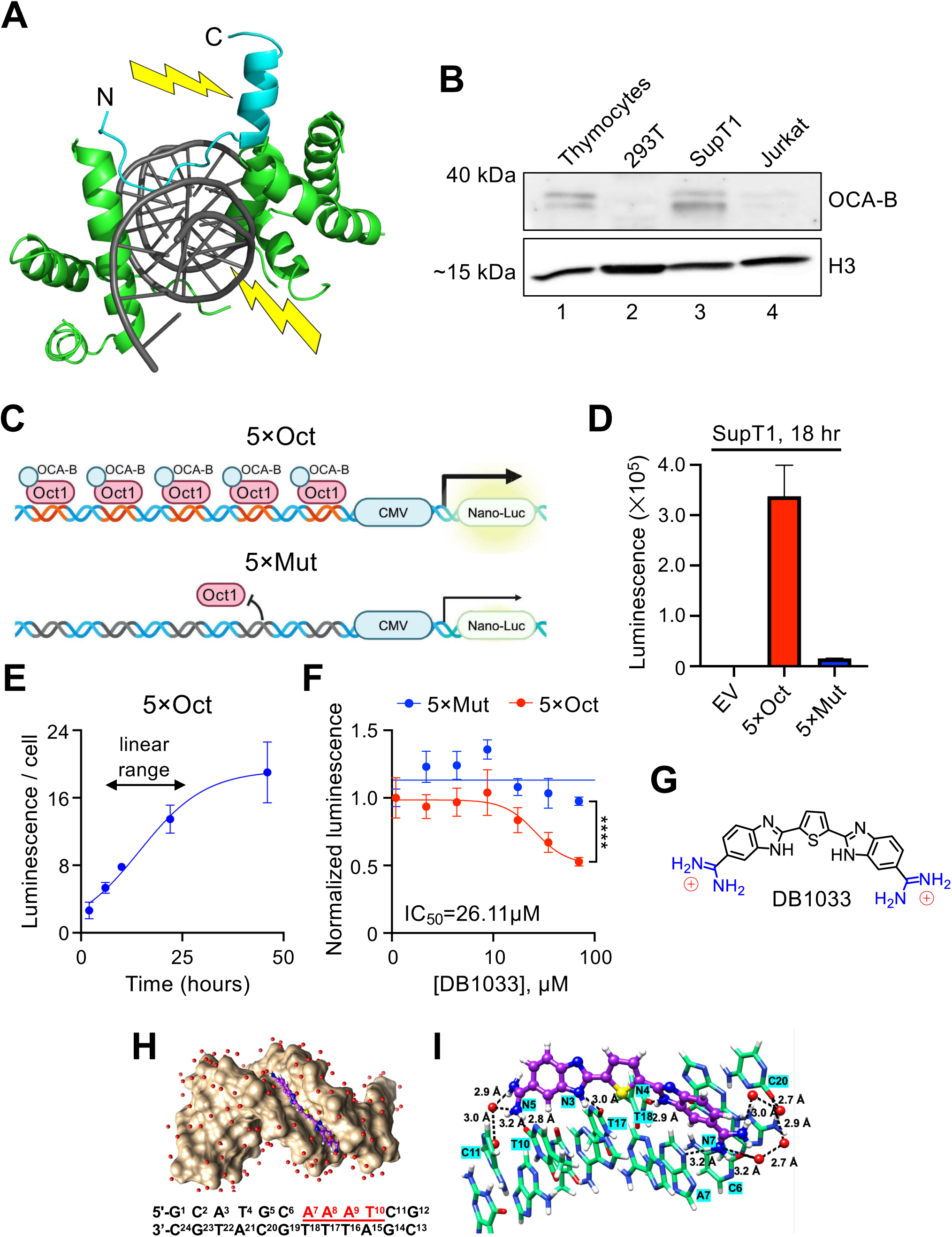
OCA-B cellular inhibition by small molecule DNA minor groove binders. (**A**) Oct1:OCA-B:DNA co-crystal structure (*45*) showing two targeting approaches: a peptide that mimics the key OCA-B alpha helix involved in downstream effector functions (top lightning bolt) (*42*), and a minor groove DNA binding molecule with potential to disrupt the unstructured Oct1 linker domain between the two DNA binding subdomains (bottom lightning bolt). (**B**) OCA-B immunoblots from SupT1 and Jurkat T cell lines (lanes 3-4). Isolated mouse primary thymocyte and 293T cell lysates were also used as positive and negative controls, respectively (lanes 1-2). Histone H3 is shown as a loading control. (**C**) Schematic for OCA-B luciferase reporter cell lines. SupT1 cells were stably transfected with reporter plasmids containing five tandem canonical Oct1:OCA-B binding sites linked to a CMV promoter upstream of secreted nLuc (5×Oct) or an identical plasmid except containing mutant binding sites used to determine nonspecific effects (5×Mut). Cells were selected with puromycin, single-cell cloned, and individual clones used subsequently. (**D**) Luciferase activity of pNL2.3 empty vector (“EV”), 5×Oct, or 5×Mut reporter lines. Cells were incubated for 18 hr after plating. n=4 biological replicate platings for each cell line. (**E**) Luciferase expression kinetics of cultured 5×Oct cells. A timecourse of 2-46 hr is shown. Averages are shown from n=5 biological replicate platings. Trendline shows logistic growth fit. (**F**) Inhibition of cellular OCA-B activity by an escalating dose of the small molecule DB1033. Values from 5×Oct and 5×Mut clones were normalized to one another in the absence of DB1033. Averages are shown across n=6 biological replicate platings. Red curve shows the “5×Oct” fit to the Hill equation. Asterisks denote results from two-tailed student T-test comparing 5×Oct and 5×Mut at the highest concentration. (**G**) Chemical structure of DB1033. (**H**) Molecular docking of DB1033 to canonical octamer B-form DNA. Red shows water molecules. DB1033 is shown in purple. Red/underlined nucleotides show the minor groove bases contacted by DB1033 docked with the lowest free energy. (**I**) Wireframe model showing DB1033 bound to the minor groove AAAT site. Hydrogen bond interactions formed between DB1033 and the minor groove AAAT site are shown with dashed lines.

To reduce the possibility of off-target hits, we tested DB1033 compound activity using an orthogonal cell-based method. We developed a nano-bioluminescence resonance energy transfer (NanoBRET) assay that measures OCA-B’s interaction with its cognate transcription factor Oct1. In this assay, 293T cells are co-transfected with an OCA-B construct carrying a HaloTag and an Oct1 construct carrying an nLuc tag (**Fig. 6A**). DB1033 again showed dose-dependent inhibition (**Fig. 6B**), indicating that DB1033 disrupts the interaction between Oct1 and OCA-B. To test this prediction at a mechanism level, we used electrophoretic mobility shift assays (EMSA) with recombinant Oct1 DNA binding domain and recombinant full-length OCA-B to measure formation of Oct1:OCA-B:DNA complexes (**Fig. 6C**). We used Cy5-conjugated double-stranded DNA to visualize complexes with cognate DNA in native polyacrylamide gels. Labeled DNA was in excess relative to protein such that changes in the ability to form a complex can be readily observed. Because of “caging effects”, changes in off-rates are not efficiently captured within a running native gel. Instead, in this assay effects of an inhibitor on complex formation are likely generated during the incubation prior to gel loading. We assembled the protein, DNA and buffer components on ice for 25 minutes in stringent conditions (150 mM KCl), and incubated the mixture for a further 5 min at room temperature ±compound prior to loading. Oct1 readily formed a complex with DNA that was augmented and slightly supershifted in the presence of OCA-B, indicating the formation of a ternary complex (**Fig. 6D**, lane 3). Inclusion of DB1033 efficiently disrupted complex formation in a dose-dependent manner (lanes 4-7).

**Fig. 6.**
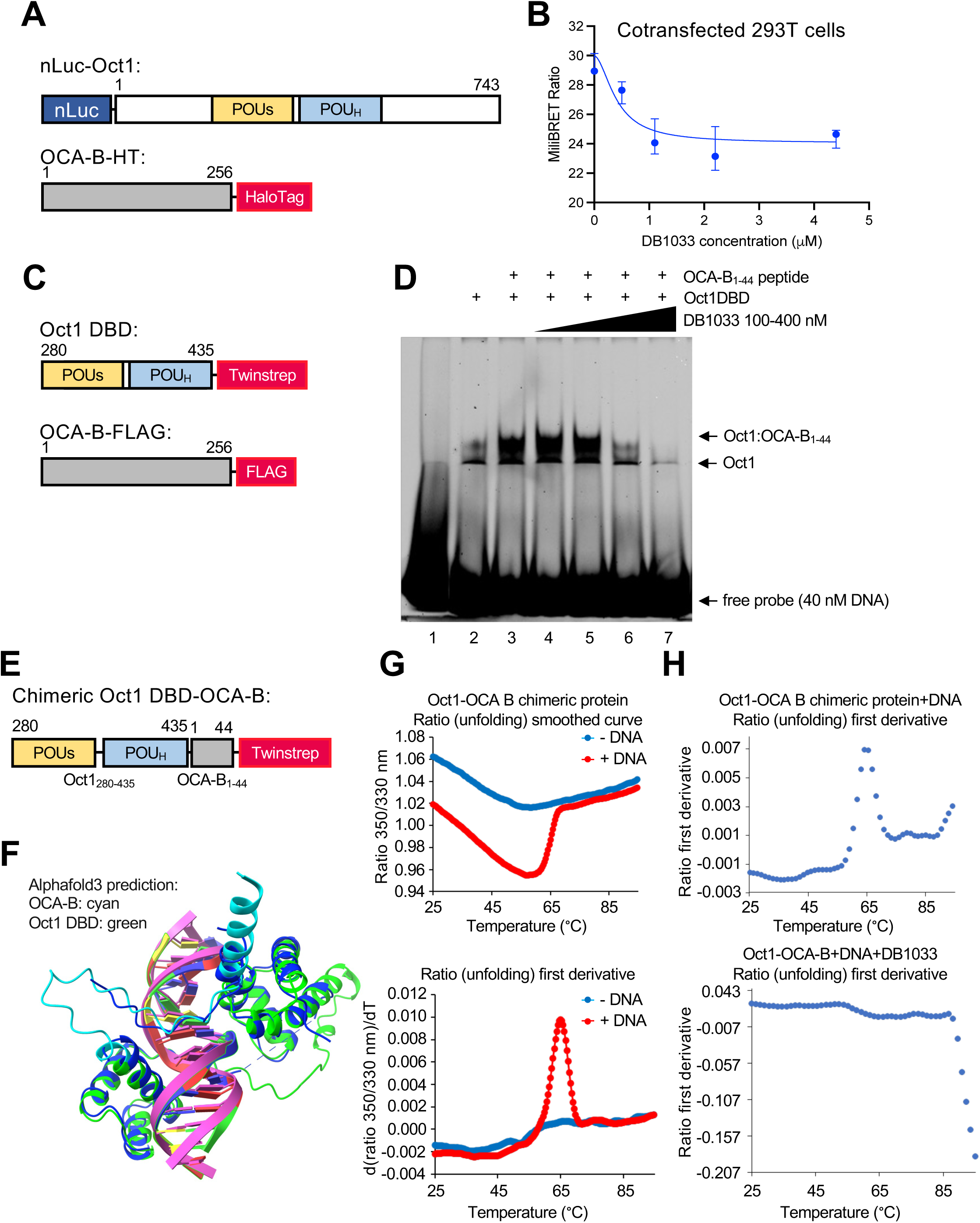
DB1033 inhibits OCA-B-Oct1-DNA complex formation. (**A**) Schematics of full-length Oct1 tagged with N-terminal nLuc and full-length OCA-B with a C-terminal HaloTag used for nanoBRET. (**B**) Quantification of nanoBRET signal obtained by co-transfecting the 293T cells with the constructs shown in A in the presence of increasing concentrations of DB1033. An average of n=3 biological replicate platings is shown. DB1033 was applied at the indicated concentrations 1 hr after seeding the cells and cells were collected after a further 24 hr.(**C**) Schematics of the C-terminally Twinstep-tagged Oct1 DNA-binding domain (Oct1_DBD_) and full-length OCA-B with a C-terminal FLAG tag used in EMSA. (**D**) Cy5-DNA scan of native PAGE EMSA showing DNA-Oct1_DBD_ complex formation with OCA-B, and concentration-dependent inhibition by DB1033. (**E**) A schematic of the Oct1_DBD_:OCA-B_1-44_ fusion protein engineered for nanoDSF. (**F**) Structure of the OCA-B_1-44_-Oct1_DBD_ complex predicted using Alphafold 3. Cyan: OCA-B_1-44_ peptide. Green: Oct1_DBD_ peptide. Dark blue shows the Oct1:DNA structure derived from X-ray crystallography (*45*). (**G**) nanoDSF measurement of the thermal stability of 10 μM Oct1_DBD_-OCA-B_1-44_ chimeric protein with or without added canonical octamer DNA. First derivative is shown below. (**H**) First derivative nanoDSF data in the presence of DNA and the presence (above) or absence (below) of 100 μM DB1033.

A complementary in vitro assay uses differential scanning fluorimetry (nanoDSF) to measure Oct1:OCA-B:DNA complex thermal stability. NanoDSF measures the gradual exposure of UV-absorbing tyrosine side chains as proteins unfold with increasing temperature. We developed a single-polypeptide chimeric molecule containing the Oct1 DNA binding domain fused to the first 44 amino acids of OCA-B, encompassing the structured part of OCA-B that contacts Oct1 (**Fig. 6E**). The first 15 amino acids of OCA-B do not appear to be well-structured and presumably serve as a linker between the two molecules. AlphaFold 3 (*55*) predicts that the chimeric molecule folds similarly to natural Oct1 and OCA-B on DNA (**Fig. 6F**). The protein was expressed in *E. coli* with N-terminal 6-His, consecutive C-terminal Twinstrep and FLAG tags. Following purification, the protein was found to be mostly pure as assessed by Coomassie staining SDS-PAGE (**fig. S7A**). Size exclusion chromatography indicated that the purified protein was mostly monodisperse, and EMSA indicated that it was able to interact with consensus octamer DNA (**fig. S7B-C**). The presence of target DNA strongly increased the stability of the chimeric protein in nanoDSF, as expected (**Fig. 6G**), while adding DB1033 in the presence of DNA destabilized the complex (**Fig. 6H**). These findings identify DB1033 as the best compound tested, indicate that DB1033 acts to disrupt Oct1:OCA-B complexes on DNA, and establish a pipeline for assessing molecules that modulate Oct1 and/or OCA-B’s interactions with one another and with DNA, their overall stability, and their transcriptional functionality.

### Small molecule Oct1:OCA-B inhibitors block α-PD-1-induced diabetes and other irAEs in NOD mice

We treated 8-week-old male NOD mice with five *i.p.* injections of 15 mg/kg DB1033 or DB2232 simultaneously with α-PD-1 antibody treatment, plus one additional dose of compound on day 11. 8-week-old male NOD.*Ocab^fl/fl^*;CD4-Cre mice were used as a positive control for protection. As with genetic OCA-B T cell knockout, DB1033 treatment completely protected mice against diabetes emergence (**Fig. 7A-B**). These results indicate 1) that DB1033 is effective in protecting mice against α-PD-1 antibody-induced diabetes, and that 2) acute inhibition at the same time as α-PD-1 antibody delivery is sufficient to protect mice against diabetes development. DB1033 toxicity was negligible or low-level, as there was a trending but statistically nonsignificant loss of weight associated with treated animals at endpoint (**Fig. 7C**). Protection was not only confined to the pancreatic islets but was also noted with salivary and lacrimal gland tissues, where leukocyte infiltration was strongly reduced (**Fig. 7D-E**). Importantly, the structurally related DB2232 compound, which was inferior in cell-based assays, was also functionally inferior at preventing diabetes (**fig. S7D-F**). These findings provide evidence of a structure-activity relationship linking compound activity to OCA-B complex inhibition. In summary, the findings show firstly that OCA-B promotes the development of irAEs in ICI-treated NOD mice while being dispensable for anti-tumor immunity, and secondly that acute small molecule inhibitor administration also blocks irAE generation.

**Fig. 7.**
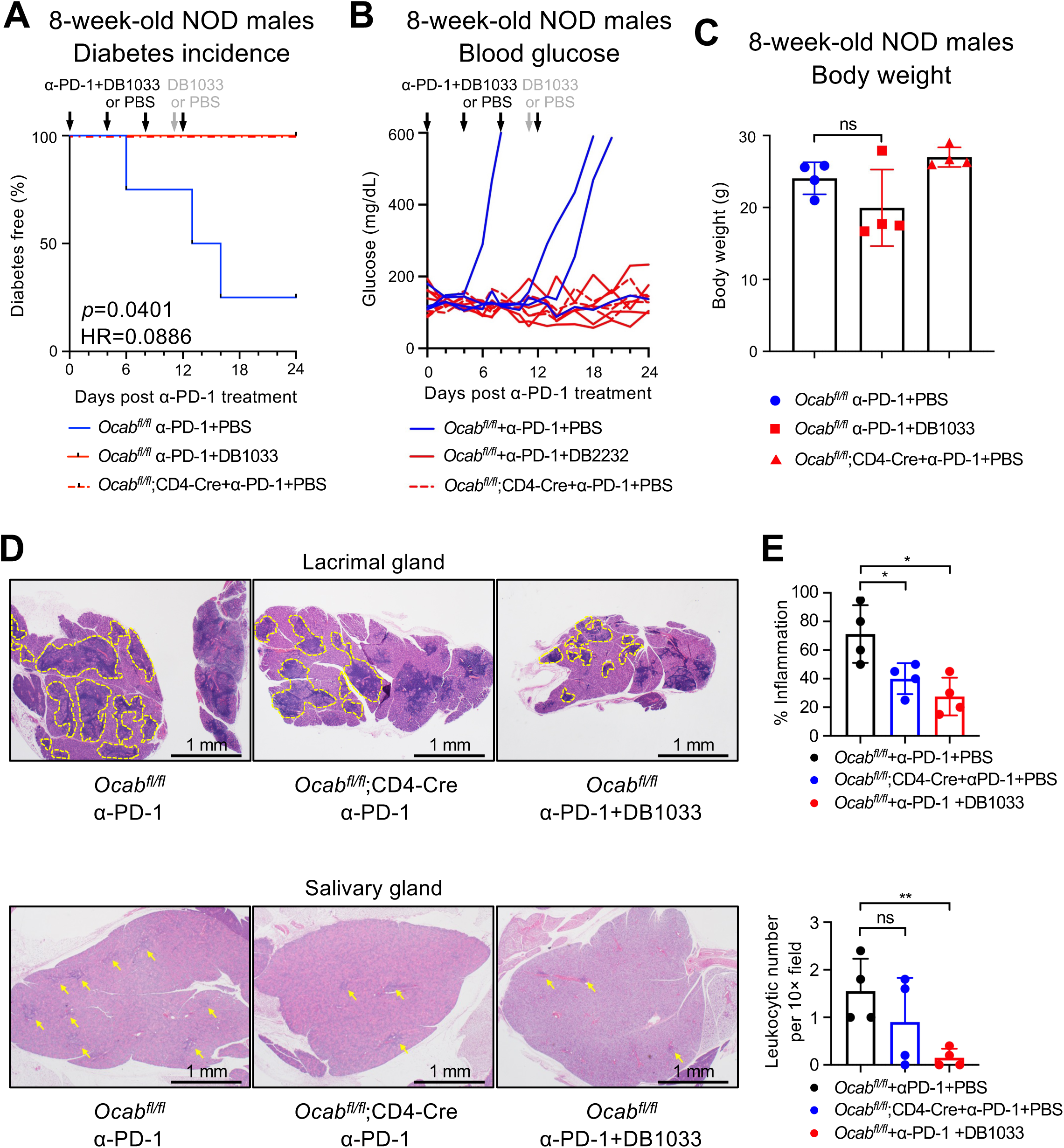
DB1033 inhibits PD-1 blockade-induced diabetes in NOD mice. (**A**) Kaplan-Meier plot of diabetes-free mice in 8-week-old male NOD.*Ocab^fl/fl^* treated with PBS (n=4), NOD.*Ocab^fl/fl^* treated with DB1033 (n=4) and NOD.*Ocab^fl/fl^*;CD4-Cre mice treated with PBS (n=4). Black arrows: 250 µg α-PD-1 and 15 mg/kg *i.p.* injections on day 0, 4, 8, 12. Grey arrow: an additional *i.p.* injection of 15 mg/kg DB1033 was administered on day 11. (**B**) Average blood glucose levels of mice in (A). (**C**) Mouse body weight on day 24 of the experiment described in (A). (**D**) Example H&E images of lacrimal and salivary gland sections harvested from 8-week-old male NOD mice injected with α-PD-1 antibodies in (A). (**E**) Quantification of the histological scoring described in (D). NOD.*Ocab^fl/fl^* treated with PBS (n=4), NOD.*Ocab^fl/fl^* treated with DB1033 (n=4) and NOD.*Ocab^fl/fl^*;CD4-Cre mice treated with PBS (n=4). Histological scoring was conducted by individuals blinded to treatment group.

## DISCUSSION

Here we show that OCA-B expression in T cells promotes irAEs following ICI treatment while minimally affecting anti-tumor responses. We also, for the first time, show that a small molecule Oct1:OCA-B inhibitor phenocopies the protection against irAEs observed using the OCA-B T cell knockouts. The latter findings indicate that acute OCA-B inhibition is sufficient to protect against irAEs and suggest the existence of a therapeutic window in which OCA-B pharmacological targeting can be used to block autoimmunity while sparing baseline immune function, including immunity to tumors.

OCA-B is one of three tissue-restricted transcription coregulators known as OCA proteins: OCA-B, OCA-T1 and OCA-T2 (*56*). Like the other two proteins OCA-B docks with the sequence-specific DNA binding POU transcription factors to regulate transcription (*43–45*). The OCA-Ts are master regulators of tuft cells, and a therapeutic vulnerability in small cell lung cancers originating from this cell type (*56*). OCA-B has important B cell-intrinsic functions including in the formation of germinal centers (*57, 58*). In T cells, OCA-B docks with the POU factor Oct1 to regulate target genes encoding cytokines and other immunomodulatory proteins (*33*). Because OCA-B expression is 50-100-fold higher in B cells compared to T cells (*31*), a calibrated dose of competitive inhibitor could selectively block T cell functions while sparing B cell-intrinsic OCA-B functions. For all three proteins, whole-body null mice are healthy, viable and fertile (*56, 59, 60*), suggesting that they a good candidate drug targets, however no small molecules targeting OCA proteins have been described.

Multiple lines of evidence show that pathogenic human and mouse progenitor-like T cells promote and sustain autoimmunity including T1D (*51, 61–63*). In mice, OCA-B promotes pathogenesis in experimental autoimmune encephalomyelitis (EAE), a model of multiple sclerosis, through maturation of stem-like cells into encephalitogenic effectors (*34*). These properties are T cell intrinsic. In a relapsing-remitting NOD.EAE model, OCA-B T cell-deficiency specifically protects mice from relapse (*34*). On the NOD background, both OCA-B whole body and T cell-specific knockout block the onset of spontaneous T1D (*42*). Using more aggressive T1D mouse models including CD25-mediated T_reg_ depletion and the CD8-restricted, autoreactive NY8.3 transgenic line revealed partial protection, while using the CD4-restricted BDC2.5 and CD8-restricted RIP-mOVA-OT-1 models showed no differences (*42*). The protection conferred by OCA-B loss is therefore strongest in more chronic polyclonal models. These findings also reveal that OCA-B T cell loss does not result in global immunosuppression, which would result in broader protection. In female NOD mice with newly arisen T1D, intravenous administration of membrane-penetrating OCA-B peptide mimics briefly reverses elevated blood glucose, as well as pancreatic islet T cell infiltration and pro-inflammatory cytokine production (*42*). In contrast to the pancreas, pLN showed no change in T cell numbers or percentages, but did show reduced T cell cytokine responses following self-peptide stimulation (*42*). However, greater than three peptide injections was toxic, with mice exhibiting ruffled skin, moribund appearance and weight loss (*42*). The Tat membrane-penetrating peptide is associated with acute toxicity that can be mitigated by spacing the dose(*64–66*).

The ability to trigger endocrine irAEs using α-PD-1 treatment in NOD mice presented an opportunity to test the OCA-B T cell loss of function model in different ways. First, to test the effect of OCA-B loss in an aggressive but physiologically relevant model of autoimmune diabetes. Second, to use the diabetes and other pathologies as models for ICI-triggered irAEs. Third, to determine the role of OCA-B in anti-tumor immunity, given the importance of finding anti-irAE treatments that do not inhibit tumor control for patients with cancer. We found that OCA-B T cell loss protected NOD mice against ICI-triggered diabetes. The effect of OCA-B loss was partially dependent on the age at which ICI was administered, with 8-week-old mice showing more complete protection compared to 12-week-old mice. This finding is consistent with a model in which OCA-B loss protects NOD mice at multiple levels, one of which spans the 8-week timepoint and is PD-1 independent, and one of which spans the 12-week timepoint and is partially PD-1 dependent. Profiling the immune cells in these mice flow cytometrically indicated significant reduction in T_EX_, and a concomitant increase in cells with a progenitor-like (SLAMF6^hi^CD39^lo^) phenotype.

In contrast to the autoimmune protection conferred by OCA-B loss, immune responses and tumor control were preserved either with or without α-PD-1 treatment. No differences were observed using immune-cold B16F10 melanoma cells, as well as immune-hot MC38 colon carcinoma cells and 9495 sarcoma cells. Similar to the preservation of tumor control, loss of OCA-B in T cells preserves normal primary response and viral control following infection with either the acute strain of lymphocytic choriomeningitis virus (LCMV) or the glial-tropic β-coronavirus JHMV (*31, 33, 34*). The normal T cell primary immune response phenotypes observed with Oct1 and OCA-B genetic deletion suggest the existence of a therapeutic window in which an OCA-B pharmaceutical could be used not just to block autoimmunity, but could be co-administered with ICIs to block irAEs while sparing immunity to viruses and tumors. A likely side effect of therapies targeting this pathway would be an inability of newly-generated memory CD4^+^ T cells to mount robust recall responses (*31, 33*). Our findings indicate that OCA-B promotes a memory-like population of CD4^+^ T cells important for immune responses to tumor re-challenge and relapse (manuscript in prep). An OCA-B inhibitor may therefore need to be withdrawn prior to any potential tumor relapse.

Despite the prevailing but erroneous idea that transcription factors are “undruggable,” there are multiple historical(*64, 67–69*), and recent(*70–84*) counterexamples, the latter enabled by technologies such as PROTAC(*85, 86*). Oct1’s wide expression pattern and role in somatic stem cell function (*87, 88*) make it less attractive as a drug target relative to OCA-B, which is highly tissue-specific. Whole-body OCA-B knockouts are viable and fertile (*59, 60*). We show that systemic administration of a small molecule inhibitor, DB1033, into NOD mice blocks the emergence of α-PD-1-induced diabetes. These molecules disrupt Oct1:OCA-B:DNA complexes in cell culture and in vitro. The finding that DB1033 administration concomitant with α-PD-1 antibody treatment is sufficient to block the induction of diabetes and other autoimmune pathologies suggests that OCA-B actions close to the time of injection, and therefore spatially proximate to the pLNs and pancreatic islets, rather than development of auto-reactive T cells in an OCA-B deficient environment from the double-positive thymocyte stage onwards, mediates autoimmune protection. Moreover, we show that a structurally related compound, DB2232, that is inferior in attenuating OCA-B-dependent transcription in cell culture and in disrupting Oct1:OCA-B complexes in vitro is also functionally inferior at protecting NOD mice against ICI-triggered diabetes, strongly suggesting that these compounds act through OCA-B disruption. These reagents provide a conceptual milestone as a first-in-class small molecule Oct1/OCA-B inhibitor that can reduce autoimmunity. More work is necessary to develop superior and more drug-like small molecule inhibitors.

## MATERIALS AND METHODS

### Design

The objective of the study was to determine the effect of OCA-B deletion in mouse T cells on autoimmune diabetes induced by ICI treatment and on immunological control of tumors in the absence and presence of ICIs, as well as to test the abilities of small molecule inhibitors. Methods included spatial transcriptomics to assess human *POU2AF1* in the context of T1D, assessment of glucose levels and tumor size, flow cytometry and histology to assess degrees of infiltration and the qualitative nature of the cells, and transcriptional reporter, nanoBRET, EMSA and nanoDSF measurements of small molecule efficacy. Sample sizes (n=3–12) were determined to be sufficient for the type of assay and statistical analysis. Excepting histological scoring, investigators were not blinded when conducting or analyzing the experiments outlined in this study. Mice were randomly assigned to treatment groups and repeat experiments were performed in both males and females where appropriate (e.g., where tumor engraftment was feasible), with no apparent sex differences.

### Spatial transcriptomics

Imaging-based spatial transcriptomics of pancreatic tissue was prepared using the network of Pancreatic Organ Donors with Diabetes (nPOD) donor tissue and the Xenium Prime in situ gene expression platform (10X Genomics) according to the manufacturers protocol. Pancreas sections were obtained from the nPOD donor 6551 (RRID: SCR+014641; https://npod.org/), a 20 year-old male with early (6 months onset) T1D (AAb count: 4) with approval from the University of Florida Institutional Review Board (IRB #201600029). Output data were processed as previously described (*46*). Xenium data were visualized using Xenium Explorer 4 software (v4.0.0).

### Laboratory mice

All mice used in this study were non-obese diabetic (NOD) background, with the exception of B16-F10 and MC38 tumor injections which used C57BL/6 mice. *Pou2af1* (*Ocab*) conditional (floxed) mice on the C57BL/6 and NOD backgrounds were generated as described previously (*31, 42*). All animal experiments were approved by the University of Utah Institutional Animal Care and Use Committee (IACUC approval 00001553).

### α-PD-1-induced diabetes

250 µg α-PD-1 (clone RMP1-14, Cat# BE0146, Bio X Cell) or rat IgG2a isotype control (Cat# BE0089, Bio X Cell) was intraperitoneally (*i.p.*) administered into NOD.*Ocab^fl/fl^* or *Ocab^fl/fl^*;CD4-Cre every 4 days as indicated. Antibody was diluted in PBS for injection of 200 µL per mouse.

### Histology

Salivary glands and lacrimal glands were isolated and fixed in 10% formalin. After 24-72 hr, tissues were transferred to 70% ethanol and embedded in paraffin, sectioned at 5∼7 μm though the middle of the tissue along the long axis, and H&E staining was performed. Slides were examined and imaged using an Olympus BX53 microscope with UPlanFL objectives, Olympus DP73 camera, and Olympus cellSens imaging software. For lacrimal glands, inflammation was assessed by quantifying the percentage of the gland area with inflammation. For salivary glands, leukocytic number per 10× field was quantified by counting the numbers of inflammatory foci and dividing by the number of 10× fields of the whole tissue.

### Secondary lymphoid organ and tumor harvest

Spleens and lymph nodes were ground and filtered through a 70 µm nylon cell strainer (VWR #76327-100). Red blood cells were removed by Ammonium-Chloride-Potassium (ACK) lysis buffer (10 mM KHCO_3_, 150 mM NH_4_Cl, and 0.1 mM Na_2_EDTA). Immune cells were recovered from MC38 tumors using a published method (*89*) in which tumors were manually disrupted by grinding between the frosted end of slide glass in a 6-well plate with 3 mL of PBS plus 4% FBS, and filtered through a 40 µm nylon cell strainer (VWR #76327-098). 9495 tumors were mechanically disrupted in 6-well-plate and digest in RPMI media containing 16mg/mL collagenase D (Sigma-Aldrich #11088858001) and 10mg/mL Dnase (Sigma-Aldrich #10104159001) at 37°C in CO_2_ incubator for 45 minutes and filtered through a 40 µm nylon cell strainer, spined in 37.5% percoll gradient, and red blood cells (RBCs) were lysed to produce single-cell suspensions.

### Pancreatic islet harvest

Lymphocytes were isolated based on a protocol from Jing et al. (*90*). Pancreata were harvested and perfused with collagenase 4 (Worthington #LS004188) at 37°C, and islets were picked using a 10 µL pipette. Islet single-cell suspensions were obtained by vortexing in cell dissociation buffer (ThermoFisher Scientific #13151014).

### Flow cytometry

Single-cell samples were stained with antibodies (1:100 diluted in FACS buffer) for flow cytometric analysis. For intracellular staining, cells were cultured in RPMI1640 (ThermoFisher Scientific #11875093) supplemented with 10% FBS, 1% penicillin/streptomycin (ThermoFisher Scientific #15140122) and 1% GlutaMAX (ThermoFisher Scientific #35050061), 1 µL/mL Brefeldin A (BD Biosciences #555029), 50 ng/mL phorbol 12-myristate 13-acetate (Sigma-Aldrich #P1585) and 1 ug/mL ionomycin (Sigma-Aldrich #I0634) for 4 hr. Cells were stained for surface markers and fixed using Cytofix/Cytoperm (BD Biosciences #554714) following the manufacture’s protocol, and resuspended in Perm/Wash buffer for intracellular cytokine staining. For intranuclear staining, cells were stained for surface markers, fixed using the Foxp3/Transcription Factor Staining Buffer Kit (ThermoFisher Scientific #00-5523-00) and stained for intranuclear markers. Flow cytometry was performed using a BD Fortessa (BD Biosciences). Data analysis was performed using FlowJo software (BD Biosciences).

### Cell lines

Mouse MC38 cell line was a gift from Ryan O’Connell (University of Utah). B16-F10 cells were a gift from the lab of Matthew A. Williams (University of Utah). MC38 cells were grown in DMEM (ThermoFisher Scientific #11965092) supplemented with 10% FBS, 1% penicillin/streptomycin, 1% MEM-NEAA (ThermoFisher Scientific #11140050) and 1% HEPES-Cl pH 7.3 (ThermoFisher Scientific #15630130). B16-F10 cells were grown in DMEM supplemented with 10% FBS and 1% penicillin/streptomycin. NOD 9495 cells were grown similar to a prior report (*54*) in DMEM (no glutamine, no sodium pyruvate, ThermoFisher Scientific #11960044) supplemented with 10% FBS (ThermoFisher Scientific #A5256701) 1% penicillin/streptomycin, 1% sodium pyruvate, 1% 100× MEM-NEAA, 1% GlutaMAX (ThermoFisher Scientific #35050061). SupT1 human T lymphoblast cells were grown in RPMI1640 supplemented with 5% FBS, 5% calf serum (Sigma), 1% penicillin/streptomycin and 1% GlutaMAX at 37°C with 5% CO_2_.

### Tumor cell line engraftment

All tumor experiments used 6- to 10-week-old C57BL/6 or 6-8-week-old NOD background mice. For injections of B16-F10 and MC38, after trypsinization, cultured tumor cells were resuspended in PBS and 1×10^6^ and 5×10^5^ tumor cells were *s.c*. injected into the hind flank of mice, respectively. MC38 tumor-bearing mice were treated with 300 µg α-PD-1 or PBS control. For the injection of NOD 9495 tumor line, after trypsinization, cultured tumor cells were resuspended in PBS and 1.5×10^6^ tumor cells were *s.c*. injected into the hind flank of mice as described (*54*). To treat tumor-bearing NOD mice while minimizing diabetes induction, 200 µg α-PD-1 or PBS control was used. To minimize complications from hyperglycemia, mice were administered insulin pellets when blood glucose levels reached 2250 mg/dL for two consecutive days.

### Small molecules

Synthesis of DB1033, DB2232, DB2658, DB2379, DB2559, and DB2385 has been previously reported (*91*). All compounds were dissolved in 100% DMSO and stored at 5mM at −80°C for later use. For cell-based assays, stock solutions or DMSO vehicle controls were diluted in culture media. For EMSA, compounds were diluted in 0.6X buffer D (*92, 93*). For mouse injections, sterile PBS was used to dissolve the compounds.

### Generation of SupT1 luciferase reporter cell lines

Reporter constructs were generated using a modified secreted nano-luciferase vector, pNL2.3 (Promega #N108A), with the hygromycin resistance cassette replaced with puromycin resistance cassette (to generate pNL2.3-Puro). Gene blocks (Integrated DNA Technologies) were synthesized containing 5 canonical Oct1 binding motifs (ATGCAAAT, 5×Oct) or mutant motifs (CAAACTAC, 5×Mut) upstream of a minimal CMV promoter. Gene blocks were subcloned into the multiple cloning site of pNL2.3-Puro using EcoRV and HindIII restriction sites. To generate stable transfectants, human SupT1 cells were electroporated using Neon (ThermoFisher Scientific) with 1 μg of either the pNL2.3-Puro empty-vector (EV), pNL2.3-Puro 5×Oct (5×Oct), or pNL2.3-Puro 5×Mut (5×Mut) construct.

Electroporation parameters were: 1700 V, 10 milliseconds, ×3 pulses. Electroporated cells were allowed to recover for 2 days, then selected for stable DNA integration and expanded in complete media containing 1 μg/mL puromycin (ThermoFisher Scientific #A1113803) for 2 weeks. Cells were then single cell-cloned by limited dilution in complete media containing 1 μg/mL puromycin.

### SupT1 OCA-B activity assays

1.0×10^5^ 5×Oct and 5×Mut reporter cells were seeded into wells of 96-well plates were cultured in 100 µL complete RPMI in complete medium with serially diluted compounds for 18-20 hr. Supernatant was collected and mixed with 1:50 buffer-diluted Nano-Glo luciferase substrate (Cat# N1110, Promega). Luminescence was measured using a Modulus Microplate Multimode Reader (Promega).

### In vivo treatment of PD-1 blockade-induced diabetes

NOD.*Ocab^fl/fl^* and NOD.*Ocab^fl/fl^*;CD4-Cre mice were *i.p.* administered with α-PD-1 and 15 mg/kg of DB1033 or DB2232 was administered *i.p.* to the mice on day 0, 4, 8, and 12, and one additional dose of DB1033 or DB2232 on day 11. Mice were sacrificed on day 24.

### Construction and purification of recombinant Oct1 DNA binding domain Oct1-fusion proteins

A cDNA for the Oct1 DNA binding domain (comprising POU_S_ and POU_H_ subdomains) fused to OCA-_B1-44_ was ordered from IDT and included an N-terminal 6-His, consecutive C-terminal Twinstrep and FLAG tags. the cDNAs were cloned into the pET28a vector (Invitrogen) between the *Nco*I and *Hind*III sites. The plasmids were transformed into Rosetta 2 (DE3) pLysS competent cells (Sigma) using 50 µg/ml kanamycin as a selectable marker. Protein was expressed using 1 mM IPTG (Sigma) at 37°C for three hr. The cells were lysed in buffer containing 20 mM HEPES, pH 7.4 (Sigma), 300 mM NaCl (Sigma), 5% glycerol (Sigma) and 1 mM DTT (ThermoFisher Scientific) containing EDTA-free protease inhibitor cocktail (Sigma) at 2X concentration. Protein was bound to a Histrap column (Cytiva) using the same buffer and eluted with buffer containing 400 mM imidazole (Sigma). The protein was further purified by size exclusion chromatography on a Superdex increase 75 10/300 GL column (Cytiva.inc). The column was loaded using PBS containing 1 mM DTT. The protein was analyzed on 12% SDS-PAGE gel visualized by Coomassie staining.

### NanoBRET

The NanoBRET donor plasmid was made by cloning nLuc in-frame with the N-terminus of full-length human Oct1 between the *Nco*I and *Xba*I restriction sites of pACE-MAM2 (Geneva Biotech). The acceptor plasmid was made by cloning full-length human OCA-B containing a C-terminal HaloTag into pHTC vector (Promega) between the *Eco*RI and *Xho*I sites. nanoBRET was performed using the NanoBRET protein:protein interaction kit (Promega) following the manufacturer protocol. Briefly, 2×10^6^ 293T cells were plated in 6-well plates were transfected with a total of 2 μg of donor plus acceptor plasmid at a ratio of 1:100. After 24 hr, cells were harvested and counted. 2×10^5^ resuspended cells were seeded into the wells of 96-well plates using DMEM lacking phenol red (ThermoFisher Scientific) containing 10% FBS. After 1 hr, DB1033 or DMSO vehicle control were also added at the indicated concentrations in biological triplicate. After 24 hr, the Nanuck substrate and Halo-ligand were added before measuring BRET ratio using plate reader (Spectromax iD5, Molecular Devices Inc.). The miliBRET ratio was plotted with respect to the DB1033 concentration.

### EMSA

Published protocols (*92, 93*) were used for EMSA. The octamer sequence probe was 5’ labeled with Cy-5 and used at a final concentration of 40 nM. The probe was incubated with 10 nM Oct1-OCA B chimeric protein ±equimolar OCA B (1–44) peptide over ice for 15 min prior to the addition of 100 nM-400 nM DB1033, then further incubated over ice for another 15 min. Reaction mixtures were electrophoresed through 6% native polyacrylamide gels. Images were collected using a Molecular Dynamics Typhoon scanner using the Cy-5 fluorescence filter.

### NanoDSF

Tryptophan fluorescence at 330 nm and 350 nm were simultaneously monitored over a temperature range from 25°C to 95°C using a NanoTemper Technologies nanoDSF instrument with an excitation wavelength of 280 nm based on published procedures (*94*). The Oct1-OCA B chimeric protein complexed with double-stranded DNA containing an Oct1-OCA-B binding site was incubated on ice for 15 min. Protein lacking DNA was used as a control. The effect of DB1033 compound or DMSO control on the protein DNA complex were measured by comparing the melting point of the protein/DNA complex ±DB1033 compound. 10 µl from each sample was loaded into each capillary avoiding any air bubbles. The fluorescence intensity ratio and its first derivative were calculated with the manufacturer’s software (PR.ThermControl, version 2.1.2). The smoothened curves and the first derivative were plotted in using Microsoft Excel software. The peak from the first-order derivative indicates the melting temperature of the sample protein sample.

### Statistical analyses

Two-tailed student’s t-tests were used to determine statistical differences in experiments with pairwise comparison of data points between two groups. Two-way ANOVA analyses were used to determine statistical differences in the tumor growth curves. Survival curves used a Mantel-Haenszel test to calculate hazard ratio and a Mantel-Cox test to calculate statistical significance. All error bars represent ±standard deviations unless otherwise noted. For tumor growth assays, error bars denote ±SEM. For all other experiments, error bars denote ±standard deviation. For all figures, ns (not significant), *p* > 0.05; *, *p* ≤ 0.05; **, *p* ≤ 0.01; ***, *p* ≤ 0.001; ****, *p* ≤ 0.0001.

## Supplemental Figure Legends

**Fig. S1. OCA-B deletion in T cells protects NOD mice from spontaneous and PD-1 blockade-induced diabetes.** (**A**) Kaplan-Meier plot of diabetes-free mice in 12-week-old female littermate control NOD.*Ocab^fl/fl^* (n=5) or experimental NOD.*Ocab^fl/fl^*;CD4-Cre (n=6) mice receiving 250 µg α-PD-1 antibody injections on day 0, 4, 8, 12, and 16, indicated by black arrows. Spontaneous T1D was monitored in NOD.*Ocab^fl/fl^* (n=5) and NOD.*Ocab^fl/fl^*;CD4-Cre (n=5) mice receiving PBS. (**B**) Blood glucose levels were plotted from individual mice receiving α-PD-1 antibody in (A). (**C**) Flow cytometry plots showing percentages of CD8^+^PD-1^+^ islet T cells collected from 12-week-old mice 6 days post-α-PD-1 treatment. (**D**) Quantification of PD-1^+^ T cells among CD8^+^ T cells from islets of *Ocab^fl/fl^* (n=3) and *Ocab^fl/fl^*;CD4-Cre mice (n=5). (**E**) Flow cytometry plots showing percentages of SLAMF6^+^CD39^-^and SLAMF6^-^CD39^+^ among CD8^+^PD-1^+^ T cells in islets collected from 12-week-old mice 6 days post-α-PD-1 treatment. (**F**) Quantification of flow cytometry data in (E) from *Ocab^fl/fl^* (n=3) and *Ocab^fl/fl^*;CD4-Cre mice (n=5). Percentages of CD8^+^PD-1^+^SLAMF6^-^CD39^+^ among total islet cells were shown. (**G**) Flow cytometry plots showing percentages of SLAMF6^+^CD39^-^ and SLAMF6^-^CD39^+^ among CD4^+^PD-1^+^ T cells in islet collect from 12-week-old mice 6 days post-α-PD-1 treatment. (**H**) Quantification of flow cytometry data in (**G**) from NOD.*Ocab^fl/fl^* (n=3) and NOD.*Ocab^fl/fl^*;CD4-Cre mice (n=5).

**Fig. S2. OCA-B deletion in T cells results in the accumulation of CD62L^hi^ T_PEX_ in pancreatic islets.** (**A**) Flow cytometry plots showing percentages of T_PEX_ (SLAMF6^+^CD39^-^) and T_EX_ (SLAMF^-^CD39^+^) in islets collected from NOD 8-week-old males without α-PD-1 treatment. (**B**) Quantification of percentages of T_PEX_ (SLAMF6^+^CD39^-^) and T_EX_ (SLAMF^-^CD39^+^) among total cells from islets of *Ocab^fl/fl^*(n=3) and *Ocab^fl/fl^*;CD4-Cre mice (n=5). (**C**) Flow cytometry plots showing expressions of CD62L and CD39 from mice in Fig. 2. Cells were pre-gated on CD8^+^PD-1^+^. (**D**) Quantification of flow cytometry data for the percentage of CD62L^+^T_PEX_, conventional CD62L^-^ T_PEX_ and CD39^+^ T_EX_ in islets from NOD.*Ocab^fl/fl^* (n=7) and NOD.*Ocab^fl/fl^*;CD4-Cre mice (n=8).

**Fig. S3. OCA-B deletion in T cells leaves B16F10 tumor growth and unaltered.** Growth of *s.c*. B16F10 tumors in C57BL/6 mice sufficient (blue, *Ocab^fl/fl^*, n=3) and deficient (red, *Ocab^fl/fl^*;CD4-Cre, n=3) for T cell-expressed OCA-B. No α-PD-1 was provided.

**Fig. S4. NOD tumor additional data. (A)** Kaplan-Meier plot of another cohort of diabetes-free mice performed with the same experimental procedures as in Figure 3. *Ocab^fl/fl^* (n=14) and *Ocab^fl/fl^*;CD4-Cre (n=9). **(B-E)** Flow cytometric analysis was performed on tumors harvested from mice with similar experiment procedures described in Fig. 4A. Representative plots and quantifications of CD4^+^ and CD8^+^ effector (CD44^+^CD62L^-^) T cells, and exhausted (PD-1^+^) CD4^+^ and CD8^+^ T cells are shown.

**Fig. S5. Effect of staurosporine on luciferase-based transcription activity in SupT1 cells.** A dose curve of staurosporine is shown using both the 5×Oct and 5×Mut cell lines. Trendlines show the fit to the Hill equation.

**Fig. S6. OCA-B cellular inhibition by additional small molecule DNA minor groove binders.** (**A**) Inhibition of cellular OCA-B activity by DB2232. Blue datapoints represent control measurements of nonspecific transcription measured in 5×Mut cells. Red datapoints represent experimental measurements from the 5×Oct line. Luciferase values were normalized to 1.0 in the absence of DB2232. Trendlines show the fit to the Hill equation. (**B-H**) Similar data for additional molecules. 5×Mut cells were used for panels (F,G,H). Cytotoxicity was measured using viable cell counts for panels (B,C,D,E).

**Fig. S7. Additional nanoDSF validation data and DB2232 effect on PD-1 blockade-induced diabetes in NOD mice. (A)** Image of a 12% SDS-polyacrylamide gel loaded with the purified recombinant chimeric Oct1 DBD-OCA-B protein and stained with Coomassie blue. Molecular weight makers are shown at left as a standard. **(B)** Size exclusion chromatography elution profile using the Oct1-OCA-B chimera. **(C)** EMSA using the Oct1-OCA-B chimera and Cy5-labeled prefer octamer consensus DNA. **(D)** Kaplan-Meier plot of diabetes-free mice in 8-week-old male NOD.*Ocab^fl/fl^* mice treated with PBS (n=7), NOD.*Ocab^fl/fl^*mice treated with DB2232 (n=4) and NOD.*Ocab^fl/fl^*;CD4-Cre mice treated with PBS (n=7). Black arrows: 250 µg α-PD-1 and 15 mg/kg *i.p.* injections on day 0, 4, 8, 12. Grey arrow: an additional *i.p.* injection of 15 mg/kg DB2232 was administered on day 11. **(E)** Average blood glucose levels of mice in (D). **(F)** Body weight of mice on day 24 of the experiment described in (D).

## Acknowledgements

We thank R. O’Connell and M. Williams for MC38 and B16F10 cells, A. Young for 9495 cells, and O. Pornillos for use of nanoDSF instrumentation. We thank O. Allen, B. Dalley and the University of Utah High-Throughput Genomics Core facility. We thank J. Marvin and the University of Utah Flow Cytometry Core facility. We acknowledge the Huntsman Cancer Institute Cancer Center Support Grant (CCSG). This work was supported by grants from the National Institutes of Health/National Institute of Allergy and Infectious Diseases (R01AI162929), the Praespero Autoimmune Fund (Canada), and an internal seed grant from the Huntsman Cancer Institute Genomics, Epigenetics and Metabolism (GEM) program to DT. Additional support was provided by National Institutes of Health grant T32 DK108736 (MAM-S) and National Institutes of Health grant P01 AI042288 (TMB). This research was performed with the support of the Network for Pancreatic Organ donors with Diabetes (nPOD; RRID:SCR_014641), a collaborative type 1 diabetes research project supported by Breakthrough T1D and The Leona M. & Harry B. Helmsley Charitable Trust (Grant#3-SRA-2023-1417-S-B). The content and views expressed are the responsibility of the authors and do not necessarily reflect the official view of nPOD. Organ Procurement Organizations (OPO) partnering with nPOD to provide research resources are listed at https://npod.org/for-partners/npod-partners/.

## Author contributions

JD, AKM, MAM-S, EPH and PB performed experiments and analyzed data. AY and TMB provided key reagents and methods. KJE acquired images and analyzed data. WDW and AAF synthesized and purified DNA minor groove binding small molecules. DT conceptualized and supervised the study. All authors contributed to the writing of the manuscript.

