## Supplemental Figures and text for "OCA-B/Pou2af1 Expression in T Cells Promotes PD-1 Blockade-Induced Autoimmunity but is Dispensable for Anti-Tumor Immunity"

Supplemental material includes:

Supplemental Figs. S1-S7

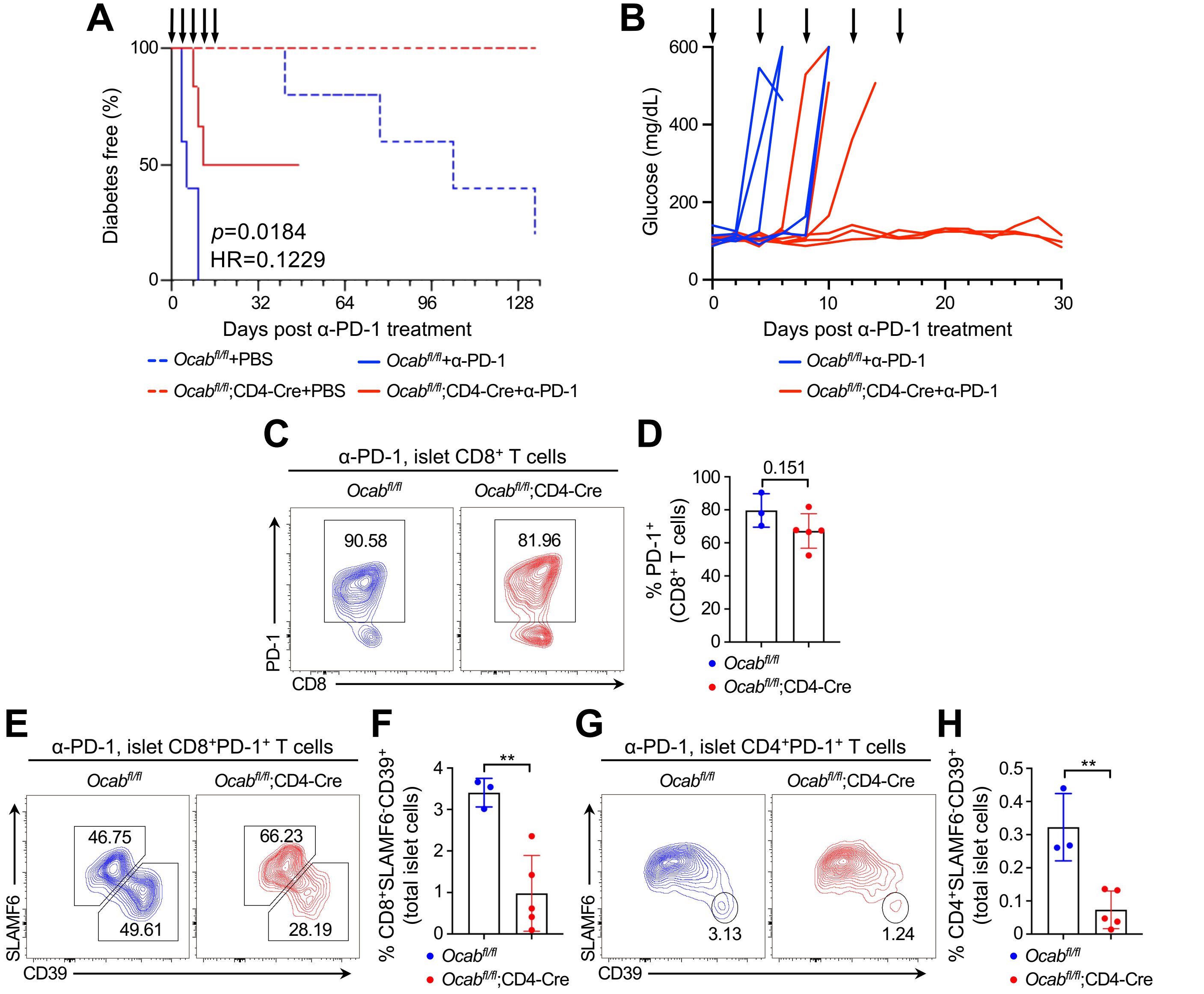


**Fig. S1. OCA-B deletion in T cells protects NOD mice from spontaneous and PD-1 blockade-induced diabetes.** (**A**) Kaplan-Meier plot of diabetes-free mice in 12-week-old female littermate control NOD.*Ocab*$^{fl/fl}$ (n=5) or experimental NOD.*Ocab*$^{fl/fl}$;CD4-Cre (n=6) mice receiving 250 μg α-PD-1 antibody injections on day 0, 4, 8, 12, and 16, indicated by black arrows. Spontaneous T1D was monitored in NOD.*Ocab*$^{fl/fl}$ (n=5) and NOD.*Ocab*$^{fl/fl}$;CD4-Cre (n=5) mice receiving PBS. (**B**) Blood glucose levels were plotted from individual mice receiving α-PD-1 antibody in (A). (**C**) Flow cytometry plots showing percentages of CD8$^{+}$PD-1$^{+}$ islet T cells collected from 12-week-old mice 6 days post-α-PD-1 treatment. (**D**) Quantification of PD-1$^{+}$ T cells among CD8$^{+}$ T cells from islets of *Ocab*$^{fl/fl}$ (n=3) and *Ocab*$^{fl/fl}$;CD4-Cre mice (n=5). (**E**) Flow cytometry plots showing percentages of SLAMF6$^{+}$CD39$^{-}$ and SLAMF6$^{-}$CD39$^{+}$ among CD8$^{+}$PD-1$^{+}$ T cells in islets collected from 12-week-old mice 6 days post-α-PD-1 treatment. (**F**) Quantification of flow cytometry data in (E) from *Ocab*$^{fl/fl}$ (n=3)

and *Ocab*$^{fl/fl}$;CD4-Cre mice (n=5). Percentages of CD8$^{+}$PD-1$^{+}$SLAMF6$^{-}$CD39$^{+}$ among total islet cells were shown. (**G**) Flow cytometry plots showing percentages of SLAMF6$^{+}$CD39$^{-}$ and SLAMF6$^{-}$CD39$^{+}$ among CD4$^{+}$PD-1$^{+}$ T cells in islet collect from 12-week-old mice 6 days post-α-PD-1 treatment. (**H**) Quantification of flow cytometry data in (**G**) from NOD.*Ocab*$^{fl/fl}$ (n=3) and NOD.*Ocab*$^{fl/fl}$;CD4-Cre mice (n=5).

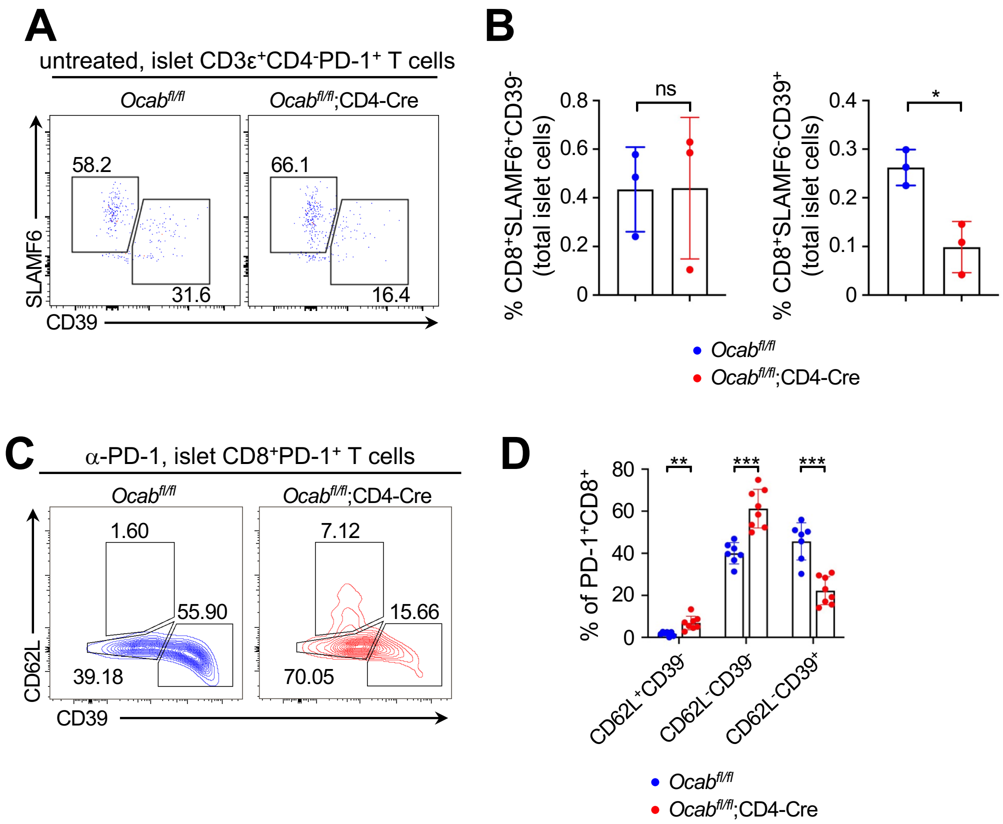


**Fig. S2. OCA-B deletion in T cells results in the accumulation of CD62L$^{hi}$ T$_{PEX}$ in pancreatic islets.** (**A**) Flow cytometry plots showing percentages of T$_{PEX}$ (SLAMF6$^+$CD39$^-$) and T$_{EX}$ (SLAMF$^-$CD39$^+$) in islets collected from NOD 8-week-old males without α-PD-1 treatment. (**B**) Quantification of percentages of T$_{PEX}$ (SLAMF6$^+$CD39$^-$) and T$_{EX}$ (SLAMF$^-$CD39$^+$) among total cells from islets of *Ocab$^{fl/fl}$* (n=3) and *Ocab$^{fl/fl}$*;CD4-Cre mice (n=5). (**C**) Flow cytometry plots showing expressions of CD62L and CD39 from mice in Fig. 2. Cells were pre-gated on CD8$^+$PD-1$^+$. (**D**) Quantification of flow cytometry data for the percentage of CD62L$^+$T$_{PEX}$, conventional CD62L$^-$ T$_{PEX}$ and CD39$^+$ T$_{EX}$ in islets from NOD.*Ocab$^{fl/fl}$* (n=7) and NOD.*Ocab$^{fl/fl}$*;CD4-Cre mice (n=8).

### Du et al. Supplemental Figure 3

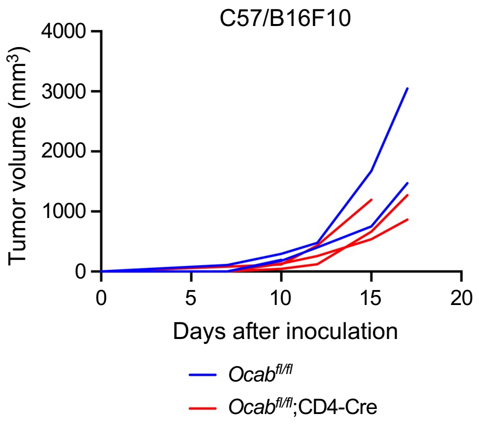


**Fig. S3. OCA-B deletion in T cells leaves B16F10 tumor growth unaltered.** Growth of *s.c.* B16F10 tumors in C57BL/6 mice sufficient (blue, *Ocab*$^{fl/fl}$, n=3) and deficient (red, *Ocab*$^{fl/fl}$;CD4-Cre, n=3) for T cell-expressed OCA-B. No α-PD-1 was provided.

### Du et al. Supplemental Figure 4

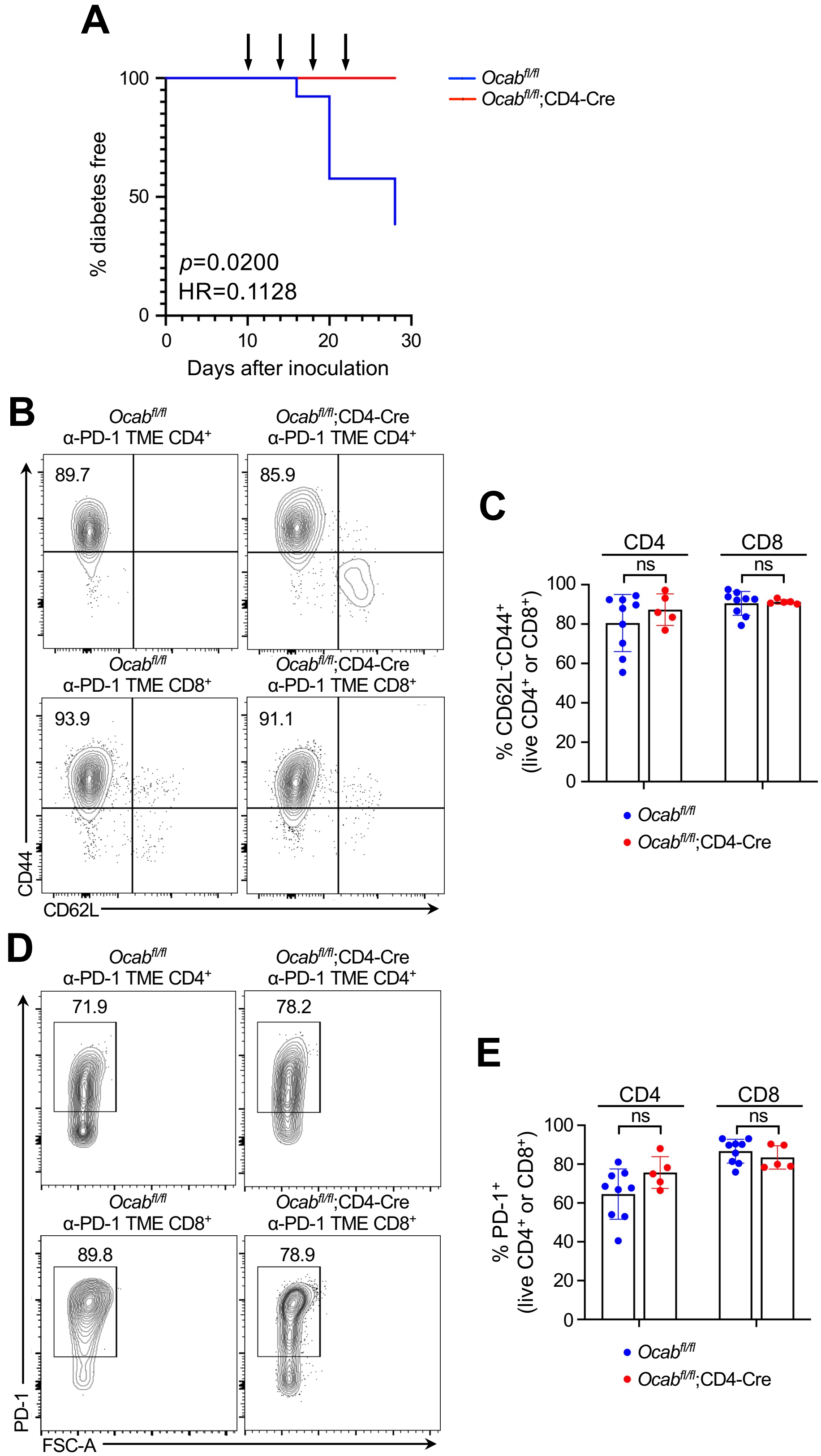

**Fig. S4. NOD tumor additional data. (A)** Kaplan-Meier plot of another cohort of diabetes-free mice performed with the same experimental procedures as in Figure 3. *Ocab*$^{fl/fl}$ (n=14) and *Ocab*$^{fl/fl}$;CD4-Cre (n=9). **(B-E)** Flow cytometric analysis was performed on tumors harvested from mice with similar experiment procedures described in Fig. 4A. Representative plots and quantifications of $CD4^+$ and $CD8^+$ effector ($CD44^+CD62L^-$) T cells, and exhausted (PD-$1^+$) $CD4^+$ and $CD8^+$ T cells are shown.

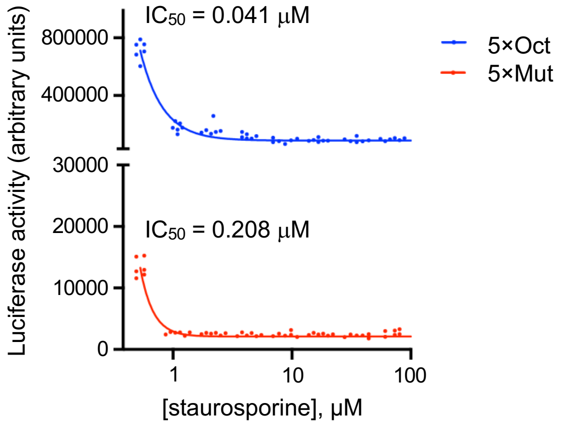


**Fig. S5. Effect of staurosporine on luciferase-based transcription activity in SupT1 cells.** A dose curve of staurosporine is shown using both the 5×Oct and 5×Mut cell lines. Trendlines show the fit to the Hill equation.

Du et al. Supplemental Figure 6

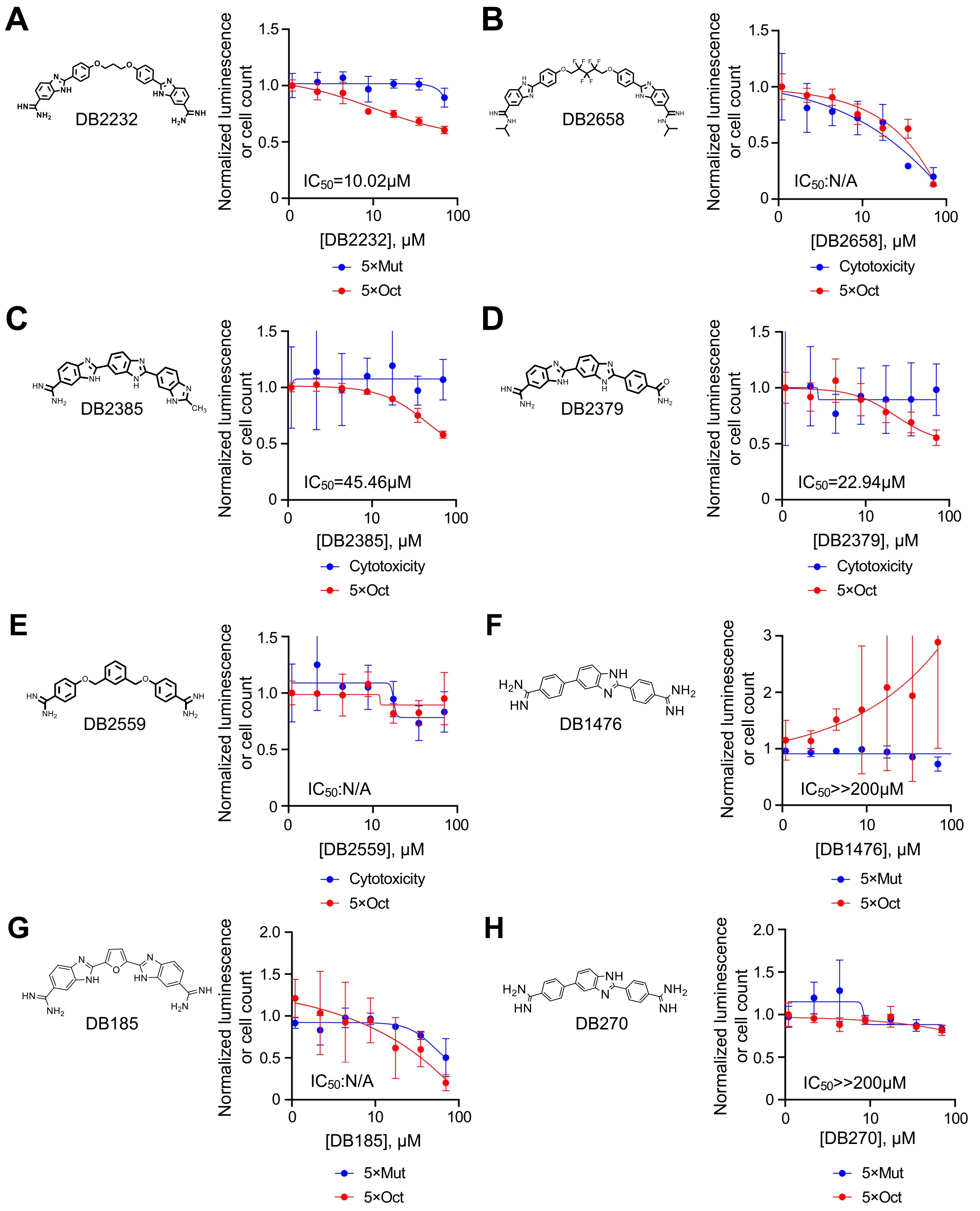

**Fig. S6. OCA-B cellular inhibition by additional small molecule DNA minor groove binders.** (**A**) Inhibition of cellular OCA-B activity by DB2232. Blue datapoints represent control measurements of nonspecific transcription measured in 5×Mut cells. Red datapoints represent experimental measurements from the 5×Oct line. Luciferase values were normalized to 1.0 in the absence of DB2232. Trendlines show the fit to the Hill equation. (**B-H**) Similar data for additional molecules. 5×Mut cells were used for panels (F,G,H). Cytotoxicity was measured using viable cell counts for panels (B,C,D,E).

### Du et al. Supplemental Figure 7

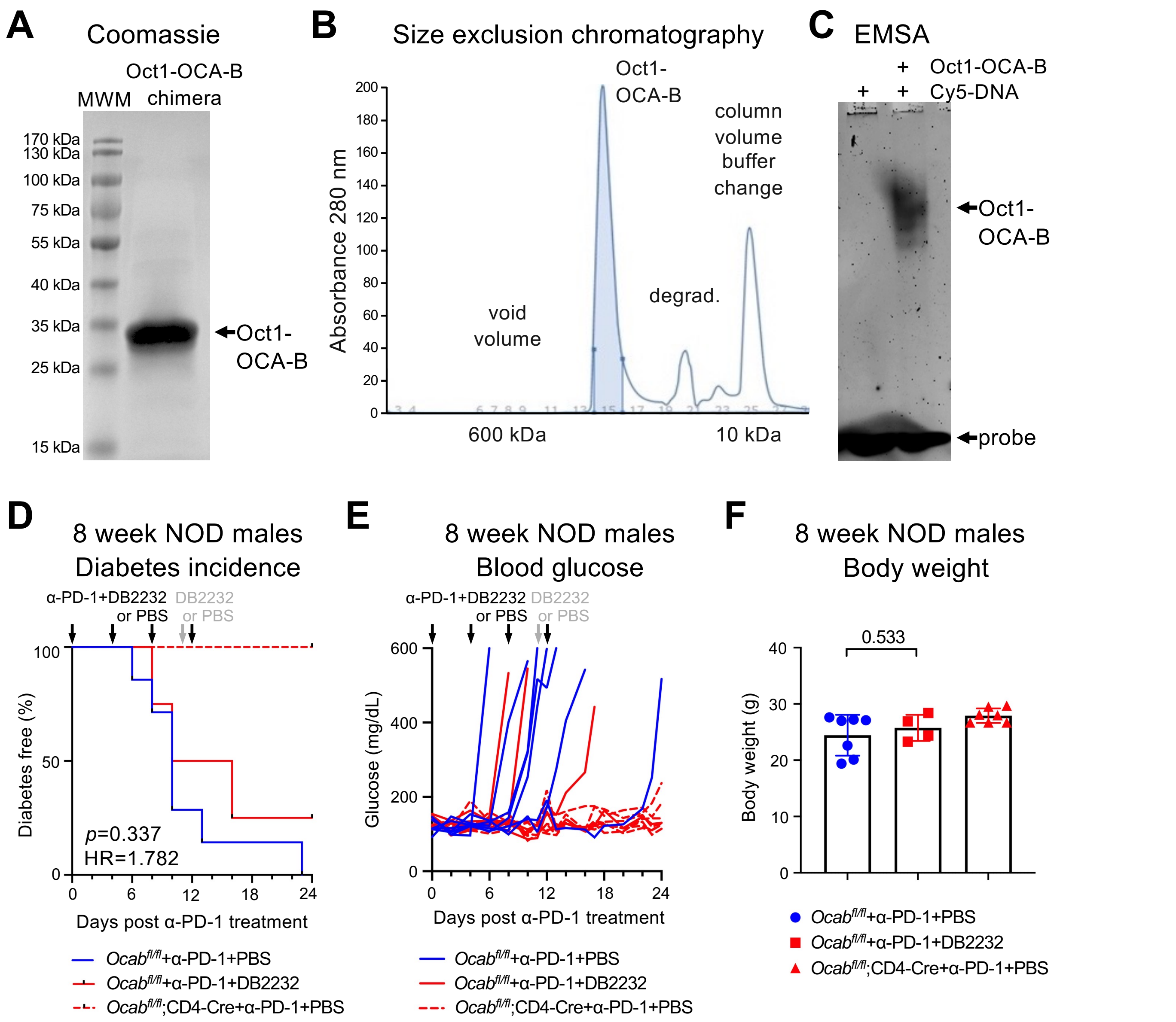


**Fig. S7. Additional nanoDSF validation data and DB2232 effect on PD-1 blockade-induced diabetes in NOD mice. (A)** Image of a 12% SDS-polyacrylamide gel loaded with the purified recombinant chimeric Oct1 DBD-OCA-B protein and stained with Coomassie blue. Molecular weight makers are shown at left as a standard. **(B)** Size exclusion chromatography elution profile using the Oct1-OCA-B chimera. **(C)** EMSA using the Oct1-OCA-B chimera and Cy5-labeled prefer octamer consensus DNA. **(D)** Kaplan-Meier plot of diabetes-free mice in 8-week-old male NOD.*Ocab$^{fl/fl}$* mice treated with PBS (n=7), NOD.*Ocab$^{fl/fl}$* mice treated with DB2232 (n=4) and NOD.*Ocab$^{fl/fl}$*;CD4-Cre mice treated with PBS (n=7). Black arrows: 250 μg α-PD-1 and 15 mg/kg *i.p.* injections on day 0, 4, 8, 12. Grey arrow: an additional *i.p.* injection of 15 mg/kg DB2232 was administered on day 11. **(E)** Average blood glucose levels of mice in (D). **(F)** Body weight of mice on day 24 of the experiment described in (D).
